# The mutation and clonality profile of genomically unstable high grade serous ovarian cancer is established early in tumor development and conserved throughout therapy resistance

**DOI:** 10.1101/2025.02.14.638365

**Authors:** Michael Diaz, Nicole Gull, Pei-Chen Peng, Kruttika Dabke, Jessica Baker, Jennifer Sun, Juan Alvarado, Benjamin Bastin, Nimisha Mazumdar, Chintda Santiskulvong, Andrew J. Li, Beth Y. Karlan, BJ Rimel, Sarah J. Parker, Simon A. Gayther, Michelle R. Jones

**Author notes:** Corresponding author:, Address: The Board of Governor’s Innovation Center, Pacific Design Center, Blue Building Suite 223, 8687 Melrose Ave., West Hollywood CA 90069. These authors contributed equally (Michael Diaz, Nicole Gull).

## Abstract

High grade serous ovarian cancer (HGSOC) is the most lethal gynecologic malignancy, killing more than 9,000 women each year in the United States alone. Nearly 80% of patients with HGSOC tumors will experience recurrence within 5 years, but little is known about the mechanisms that drive this process. Intratumor heterogeneity is believed to be a key feature of recurrence and resistance in HGSOC tumors, with early studies reporting diverse and complex mechanisms of the seeding of metastatic sites, including metastasis reseeding the primary tumor. Few studies have investigated temporal changes to clonality and structural variants through disease recurrence and the development of chemoresistance as surgical debulking is not frequently performed at this later stage. We performed multi-omic profiling in paired chemo-naive and chemoresistant tumors from 32 HGSOC patients to investigate the mutational and genomic landscape of disease progression. Somatic mutation profiles were largely conserved through disease progression. Mutational burdens did not significantly differ across recurrence but were driven by homologous recombination repair deficiency status. Clonal composition and dynamics were measured through variant allele frequency alterations as tumors progressed from primary chemo-naïve to recurrent chemo-resistant tumors. A novel candidate driver gene, *MDC1*, from the homologous recombination repair pathway, was significantly mutated and over-represented in patients with homologous recombination proficient tumors, with somatic mutations clustered in a single exon. Tumor evolution and phylogeny revealed that few changes in clonal abundance and complexity occur across the disease course in these patients. Taking structural variants into account, homologous recombination repair proficient (HRP) tumors tend to be polyclonal while homologous recombination repair deficient (HRD) tumors tend to be monoclonal, accompanied by a longer progression-free survival than the HRP patients. Three distinct classes of tumors were identified by structural variant signature analysis: tumors defined by DNA losses, tumors defined by DNA gains, and tumors defined by copy number neutral changes, which were largely defined by HRD status. Each class displayed distinct regions of the chromosome that were frequently affected by large scale SV events (>5Mb). Although no regions were frequently altered in recurrent tumors, GO analysis revealed that recurrent tumors have a significantly reduced immune response, which was not seen in the primary tumors. Ultra long read sequencing validated a majority of the SVs identified in short read sequencing and identified additional SVs undetected by short reads. These analyses identify that the phenotype of high grade serous ovarian tumors as defined by mutation and clonality profiles is established early in disease development and remain largely unchanged through chemotherapy and recurrence. This, when considered with the significant inter-patient heterogeneity identified in HGSOC, demonstrates the need for personalized therapies based on tumor profiling.

## Introduction

High grade serous ovarian cancer (HGSOC) is the deadliest subtype of the most lethal gynecologic malignancy in the US, epithelial ovarian cancer (EOC)^1^. Standard of care has remained largely the same for decades; maximal debulking surgery followed by combinatorial chemotherapy with taxane and platinum agents^2^, with the more recent addition of poly-ADP ribose inhibitor (PARPi) treatment for a subset of patients^3^. Patients initially demonstrate a strong response to platinum based treatment, however >80% cycle through disease relapse and chemotherapy and eventually develop progressive, chemoresistant tumors resulting in the patient succumbing to treatment-resistant disease^4^.

HGSOC tumors are marked by a complex mix of copy number signatures, loss of DNA repair mechanisms, tumor suppressor loss, and oncogene activation leading to highly genomically unstable tumors^5^. This genomic instability leads to marked intratumoral and intertumoral heterogeneity. Both simple somatic mutations (SSMs), such as single nucleotide variants (SNVs) and indels (insertions/deletions), and structural variants (SV) impact the epigenomic and transcriptomic profiles encoded by the tumor genome. These variants shape the phenotype of the malignant epithelial cells and other cell populations within the tumor. Previous reports that focused on SSMs have established two broad categories of primary HGSOC tumors: those with and without a loss of the homologous recombination repair pathway; termed homologous recombination repair deficient (HRD) or homologous recombination proficient (HRP)^6^. How these classifications impact recurrent tumors is not currently known.

It is hypothesized that tumors begin as a single clone that acquired a driver mutation and then progressively develops mutations leading to changes in clonal fitness. Major mutational changes lead to establishment of new clones that carry passenger mutations from their founding clone. Until recently it was unknown if the local spread of HGSOC in the peritoneum arises from monoclonal or polyclonal seeding. Recent reports indicate that both of these mechanisms occur, with examples of polyclonal and monoclonal seeding observed in different metastases from a single patient^7,8^. Previously published studies of primary tumors and/or metastases have included small cohorts and have not included matched chemo-naive and recurrent tumors from the same patient^7^. Whole exome data suggests clonal populations of chemoresistant cells develop early in tumor development, seeding metastatic sites and serving as the dormant cell population from which recurrent tumors arise, but this observation has not been confirmed in matched tumors^9^.

Previous studies using next generation sequencing (NGS) have identified high levels of copy number alterations and a high burden of SVs in HGSOC^6,10–13^. However, the recent development of direct molecule whole genome sequencing by Oxford Nanopore Technology (ONT) has facilitated the characterization of highly repetitive regions previously inaccessible by NGS, and preliminary studies have indicated these new strategies improve SV detection by up to 68%^14^. It is likely that complex high-level rearrangements have been under-characterized in previous studies, which plays a role in our poor understanding of the temporal and structural genomic features of disease progression. Here, we study a cohort of 72 matched primary and recurrent tumors utilizing a combination of sequencing methods to temporally identify and validate the heterogeneous genomic features associated with mortality and clonal evolution.

## Results

### Patient cohort

We identified a cohort of 32 HGSOC patients for whom paired chemo-naive, chemoresistant tumors, and germline DNA were available (Supplementary Table 1, Supplementary Fig 1A). All patients were diagnosed with high grade serous cystadenocarcinoma of the ovary or peritoneum at stage III or greater and received their clinical care at Cedars-Sinai Medical Center. We selected 11 patients who carry germline pathogenic variants in *BRCA1* or *BRCA2* (*BRCA1/2* carriers), the remaining patients did not carry pathogenic variants in either gene, nor any gene currently used for screening for breast and ovarian cancer. *BRCA1/2* carriers had longer overall survival, as previously shown ^15^. At the time of surgical debulking, prior to chemotherapy, germline DNA was collected from circulating blood and primary tumor samples were fresh-frozen shortly after collection^15^. At patient relapse, where tumor debulking was indicated, recurrent tumor samples were also fresh-frozen shortly after collection. These cases have extensive clinical history available, including information on the type and number of cycles of chemotherapy each patient received. Patient relapse and survival is shown in Supplementary Fig 1A and Supplementary Table 1.

### Landscape of simple somatic mutations

SSMs were called from short read whole genome sequencing (WGS) performed in epithelial enriched samples from each tumor. Five tumors lacking germline variants in *BRCA1/2* developed somatic SV or somatic point mutations in these genes (Fig 1A). Polygenic risk scores were not significantly different between *BRCA1/2* carriers and non-carriers (p=0.689; Supplementary Table 1, Fig 1A). Chromothripsis was not detected in any of the germline *BRCA1/2* carriers and 37.5% (3/8) of patients with chromothriptic tumors bore chromothripsis in both the primary and recurrent tumor. *PIK3CA* amplifications were identified in 11 tumors from 9 patients (1 *BRCA1* carrier, 1 *BRCA2* carrier, 7 non-carriers) and were far more common than *PIK3CB* amplifications, which were only found in 3 tumors. *PIK3CA* and *PIK3CB* amplifications were evenly distributed among primary and recurrences, with 6 and 5 instances respectively. We observed tumors from 15 patients with somatic loss of *PTEN*, 15 with an amplification (>3.5 copy number) of *CCNE1*, and 6 with an amplification of *RB1* (2/6 due to intergenic copy number changes) (Fig 1A and Supplementary Table 2). All tumors in our cohort contained a deleterious mutation in the *TP53* gene, however not all mutations were initially detected using automated somatic variant calling methods. Upon manual review, each mutation was identified and annotated (Supplementary Table 3). In addition to *TP53*, we identified the genes *MDC1*, *MUC22*, and *TDG* as genes commonly mutated above a background rate. In accordance with previous whole genome sequencing (WGS) profiling studies of HGSOC, there are few genes frequently mutated across the cohort ^16^.

**Fig. 1:**
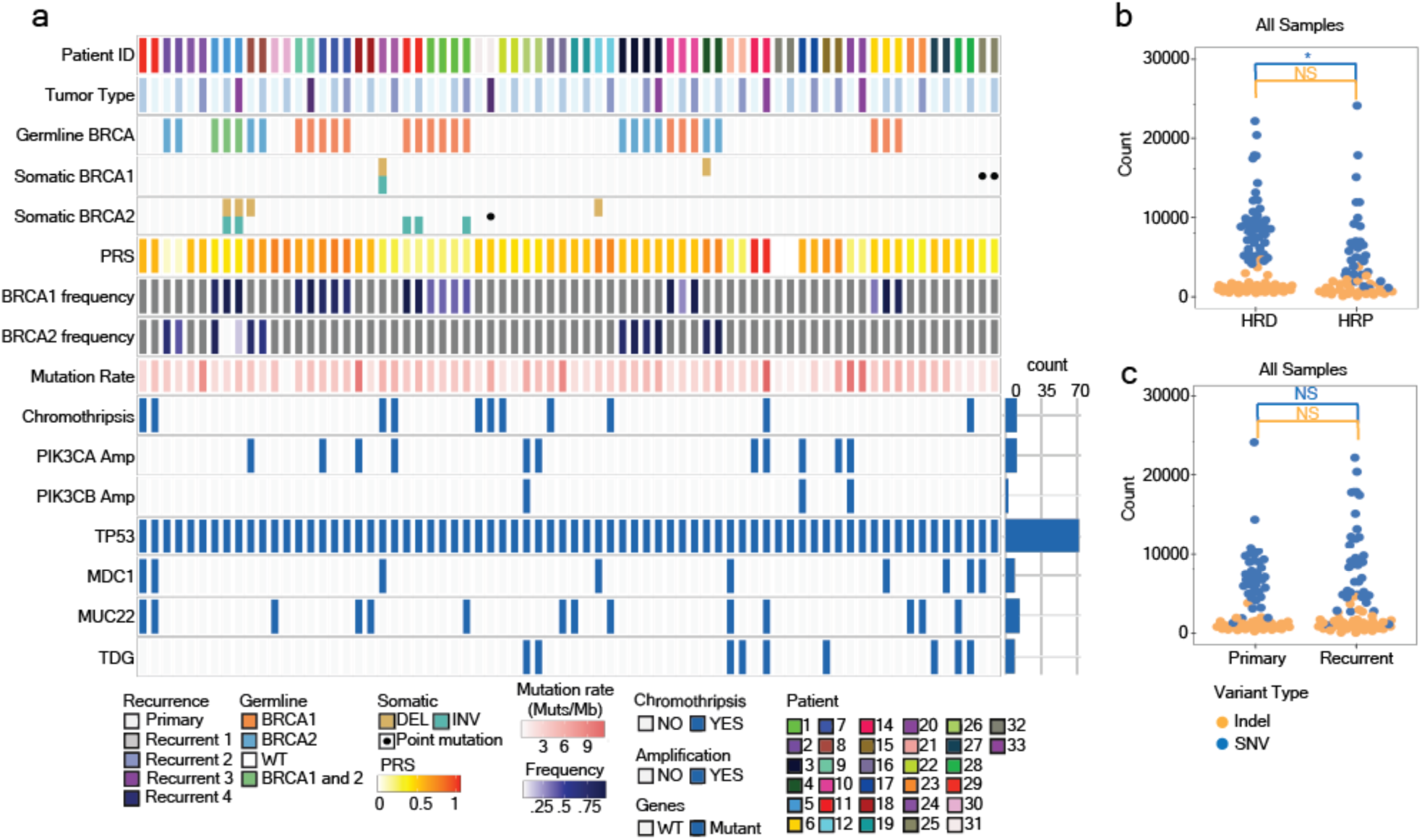
The mutational profile of HGSOC is preserved through disease recurrence. a, Descriptive graphic of tumor mutational profile, including recurrent stage, germline and somatic *BRCA1/2* variants, patient polygenic risk score (PRS), somatic *BRCA1/2* mutation frequency, mutation rate and common mutational events and commonly mutated genes within our cohort. b, Number of SNV and indel per tumor, split by HRD status. c, Number of SNV and indel per patient, split by primary or recurrent status. For b-c blue bars indicate statistics for SNV while orange bars indicate statistics for indels. *p < 0.05, NS not significant, students T test.

### Homologous recombination repair deficiency is stable throughout disease

Homologous recombination repair deficiency (HRD), as defined by germline deleterious variants or somatic mutations in *BRCA1/2*, *RAD51C*, *RAD51D*, or *BRCA1/2* promoter hypermethylation, was typically consistent across primary and recurrent tumors from the same patient. Two patients developed a new HR status throughout the disease course (Supplementary Table 2). In the fourth recurrent tumor of patient 31 (a *BRCA1/2* non-carrier), a somatic in-frame deletion with a predicted moderate effect on *BRCA1* function was observed. Of the 11 *BRCA1/2* germline carriers, reversion was observed only in the third recurrent tumor of patient 3. Patient 3 likely regained *BRCA2* function as a 36bp deletion was identified over the germline deleterious variant. This short deletion is predicted to restore the reading frame and is not predicted to be within a functional domain of the gene. Outside of these two patients, the methylation and mutation patterns of these genes were conserved. HRD status as determined by CHORD^17^, a machine learning-based method that predicts HRD status based on somatic mutation profiling, differed in some cases and did not seem to take methylation status into account. Of the 5 tumors containing *BRCA1/2* promoter methylation and the 4 tumors with *RAD51C* promoter methylation, only two were determined to be HRD by CHORD. Theoretically, as determined by CHORD, germline deleterious variants and somatic mutations in *BRCA1* did not lead to notable genomic scarring and therefore were not called as HRD for 13/22 *BRCA1* germline and somatic mutation carriers (Supplementary Table 2).

### Homologous recombination deficiency drives tumor mutational profiles

HR status, but not recurrence, had a significant effect on the number of SNVs (p=0.0167). While HRD and HRP tumors did not display significant differences in the number of indels (p=0.198), HRP tumors did have a broader distribution of indel count (Fig 1B). Recurrent tumors did not have a significantly higher SSM burden than primary tumors. However, HRD recurrent tumors have significantly more somatic SNVs than primary tumors (p=0.0314; Fig 1B, C, Supplementary Fig 1B). Copy number profiles also appeared relatively unaffected by recurrence (Figure 2A). HRD tumors have a higher count of SVs overall (p=0.00404) with significant increases in SV burden for deletions, tail to tail inversions, and translocations (p=2.72×10^−3^, 0.0387, 0.0238, 2.22×10^−3^ respectively; Fig 2B,C,D, Supplementary Table 2). There was no significant difference in the count of any individual class of SV between primary and recurrent tumors. Mean SV size across each SV class showed no difference across recurrent status (Supplementary Fig 1C, Supplementary Table 4, Fig 2E). Foldback inversions were relatively rare and not significantly affected by HRD or recurrent status (Fig 2B, C, D). The proportion of SSM and SVs shared by both the primary and recurrent tumor was not different based on HRD status (SSM HRP-25%, HRD-37% p=0.079; SV HRP-25%, HRD-34% p=0.292), however the proportion unique to the primary tumor was increased (SSM HRP-36%, HRD-29% p=.403, SV HRP-44%, HRD-35% p=0.448) (Fig 2F). There was less variability in the proportion of shared events across HRD tumors than HRP tumors (Fig 2F, Supplementary Table 5).

**Fig. 2:**
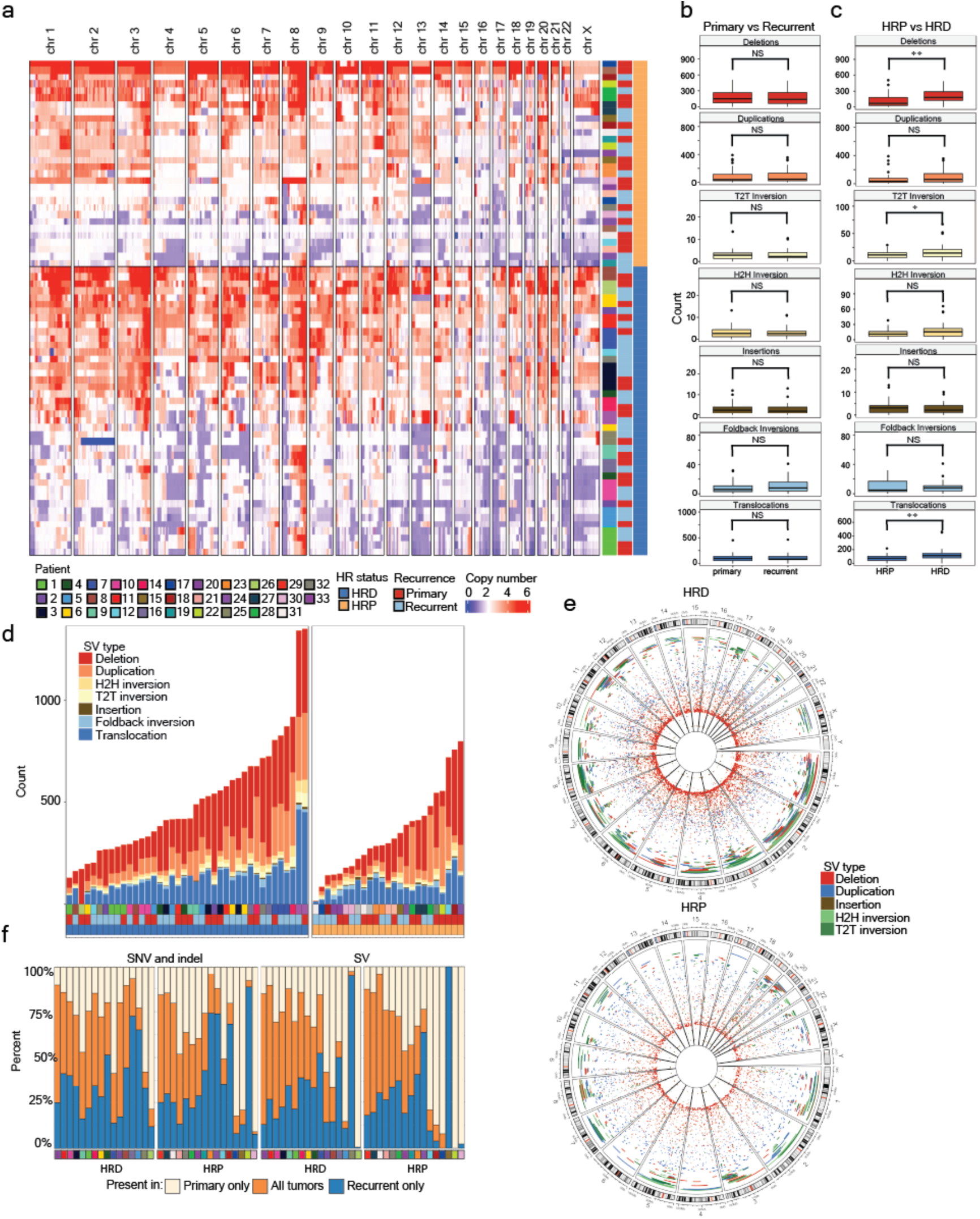
Homologous recombination status determines structural variant burden and distribution in the genome. a, Averaged copy number intervals in a 1Mb window split by HR status. Rows are clustered by copy number profiles. b, SV counts per tumor, split by primary and recurrent status. c, SV counts per tumor, split by HR status. d, Stacked bar charts depicting the total SV burden shown in c for each tumor, split by HR status. e, Circos plots depicting the location and size of each SV. The circular X axis shows location, while the Y shows the log size of the SV, relative to the chromosome in which it is shown. f, Shared and unique percentages across primary and recurrent tumors for SNV and indel (top), and SV (bottom). *p < 0.05, NS not significant, students T test.

### *MDC1* as a novel candidate driver gene in HGSOC

*MDC1* was identified as one of the most significantly mutated genes in our analysis of SSMs (Fig 1A). Nine tumors from 8 patients had SSMs in the PST region of *MDC1*, encoded by exon 11 of the *MDC1* gene, all of which were predicted to have a moderate variant effect. Five primary (2 HRD, 3 HRP) and 4 recurrent tumors (1 HRD, 3 HRP) were identified as *MDC1* somatic mutation carriers (Supplementary Fig 2). Several *MDC1* mutant tumors have regions of chromothripsis and more complex allele specific copy number profiles, irrespective of chromothripsis (Supplementary Table 2,3). *MDC1* mutants displayed a lower mutational rate than *BRCA1/2* non-carrier tumors, similar to *BRCA1/2* carrier tumors. Four of the 5 primary tumors did not have an *MDC1* mutation identified in the matched recurrent tumor. Manual review of aligned sequencing reads revealed the variant was present in all tumors from the same patient, but at a sub-clonal allele frequency that was lower than the thresholds used in our somatic variant calling pipeline (VAF<0.05). This low allele frequency may have contributed to previous studies of HGSOC not identifying this gene as being frequently mutated, as we performed somatic sequencing at an average depth of 60x. Within the Pan-Cancer Analysis of Whole Genomes (PCAWG) ovarian cancer study^18^ depth was bimodal with nodes at 38 and 60x where there was sufficient power to detect subclonal drivers that were above 30% CCF^19^. All of the *MDC1* variants identified here had a VAF between 9-30%. Mutations were confirmed using Illumina Ampliseq targeted sequencing at an average depth of 5000x. Most somatic mutations displayed a lower VAF in the Ampliseq panel than short read whole genome sequencing, and all somatic mutations in *MDC1* were detected with the second chemistry. *In silico* binding predictions failed to predict accurate structures for both *MDC1* and the interaction with one of its primary binding partners, *TP53BP1*.

We previously profiled these tumors using mass spectrometry to profile their proteome (Dabke et al, *Under Review)*, and used those data to measure MDC1 protein expression. *MDC1* demonstrated consistent protein expression across all tumor samples, with a median log2 expression of 8.38, placing it in the 8th decile when ranked by protein abundance. *MDC1* was not differentially expressed at the transcript or protein level between primary and recurrent tumors or between HRD and HRP tumors. However, it was distinctly upregulated in tumor tissues when contrasted with tumor-adjacent stromal samples, which we replicated using publicly available proteomics data sets^20^ (log2FC = 0.72; FDR = 5.27E-07)(Supplementary Table 6).

### Homologous recombination status drives SV distribution

Enrichment of SVs in hotspots across the genome was also influenced by HR status. HRP and HRD tumors displayed at least 5% of large SVs in chromosomes 1, 2, 3, 5, 7, and 8 although HRP samples possessed a greater abundance in chromosomes 5, 7, 19, 20, and 22 (Fig 2E). The largest disparity between groups in relative SV abundance was found on chromosome 19, in which 7.4% of all large SV were found in HRP tumors versus 1.2% in HRD tumors (Supplementary Table 7, Supplementary Fig 3A, 2B). Copy number profiles from chromosomes 19 and 20 also display extreme gains and losses within the same tumor, particularly in the HRP tumors (Fig 2A, Supplementary Fig 3C). HRP tumors tend to accumulate fewer SV >10Mb but have a high concentration of large SV in chromosomes 19 and 20 where 14% of all large SVs were found, with at least one variant greater than 10Mb found in 17/30 tumors. By contrast, the HRD tumors tend to have large, evenly distributed SVs that are relatively absent from chromosomes 19 and 20, where 2.7% of all large SVs were found (Fig 2E, Supplementary Table 7, Supplementary Fig 3A, B). Few regions are commonly impacted by SVs in the HRD tumors, however the q arm of chromosome 17 does contain a high proportion of SV which are frequently head to head inversions. The abundance of SVs was not unique to a particular patient’s tumors, as nearly 35% of all HRD tumors (15/42) possessed an SV of at least 1MB within chromosome 17 (Fig 2E, Supplementary Table 7, Supplementary Fig 3B). SV locations were annotated with the Gencode v34 ^21^ to determine the genes affected by each SV. GO analysis applied to the genes affected by deleterious SVs (deletion/inversion/duplication within an exon) that were shorter than 5Mb and appearing in at least 4 tumors (the most stringent filter that will produce GO results) revealed that SVs in HRP tumors were enriched in genes from lipid metabolism pathways (p=9.83×10^−4^) while HRD tumors had SVs enriched in genes in the cell adhesion pathway (p=1.24×10^− 6^)(Supplementary Fig 4A,B, Supplementary Table 8).

Fewer changes were observed in the size and distribution of SV across recurrence. (Supplementary Fig 3A, D). SV counts unique to recurrent tumors were not significantly more abundant (p = 0.661) or different in size (for size groups 1-100bp, 100bp-1kb, 1kb-5Mb, 5Mb-10Mb, and >10Mb, p = 0.564, 0.829, 0.624, 0.408, 0.884 respectively; Supplementary Table 9). Recurrent tumors did bear a high percent of very large SVs in chr 3 (17.4% of total), 61% of which were deletions. The most commonly deleted gene in this region was *FHIT*, which was deleted in 18 tumors. Loss of *FHIT* is known to induce replication stress through increased DNA damage^22^. Other frequently deleted genes include *PTPRG*, which is frequently deleted in renal cell carcinoma^23^, and *FRMD4B*, which encodes a scaffold protein.

Another marked difference in SV distribution between primary and recurrent tumors was observed in chr 17q where large SVs were mostly identified in primary tumors, not recurrent. Primary tumors were more likely to harbor large SV in chromosomes 7, 9, 10, and 17, while recurrent tumors were more likely to harbor large SVs in chromosomes 3, 4, 12, 19, and 20 (Supplementary Fig 3A, D). GO was applied to the genes affected by deleterious SVs in recurrent samples found in at least 5 patients (the most stringent filter that will produce GO results), which revealed that recurrent samples have an enrichment of SVs affecting genes from immune response pathways, driven by the two most commonly affected genes (*SLA2* and *PTK2*). This was not observed in the primary samples. The primary tumors had fewer genes deleteriously affected by SVs. Instead, genes deleteriously affected by SVs found in at least two primary tumors were subtracted from those found in at least two recurrent tumors and the resulting analysis revealed that recurrent samples displayed enrichment of SVs in genes from protein metabolism pathways (Supplementary Fig 4C, Supplementary Table 8).

### Clonal evolution occurs early in disease development

Clonal abundance (Fig 3A) changed very little through the disease course in most patients, even after primary debulking surgery followed by various lines of chemotherapy (Fig 3B, C). Clonal proportions remained relatively stable; new clones were rarely identified in the recurrent tumors, with the same clonal structure persisting through treatment in most patients (19/27; Fig 3C). In a majority of cases (18/27), the clone that comprised the greatest abundance in the primary tumor continued to comprise the greatest abundance in the recurrent tumor(s), while only 7 patients saw elimination of even a single clone during the course of chemotherapy (Fig 3C, Supplementary Fig 5). These trends were consistent across both HRP and HRD tumors, with no correlation seen between the number of SSMs and clonal burdens (Supplementary Fig 6A).

**Fig. 3:**
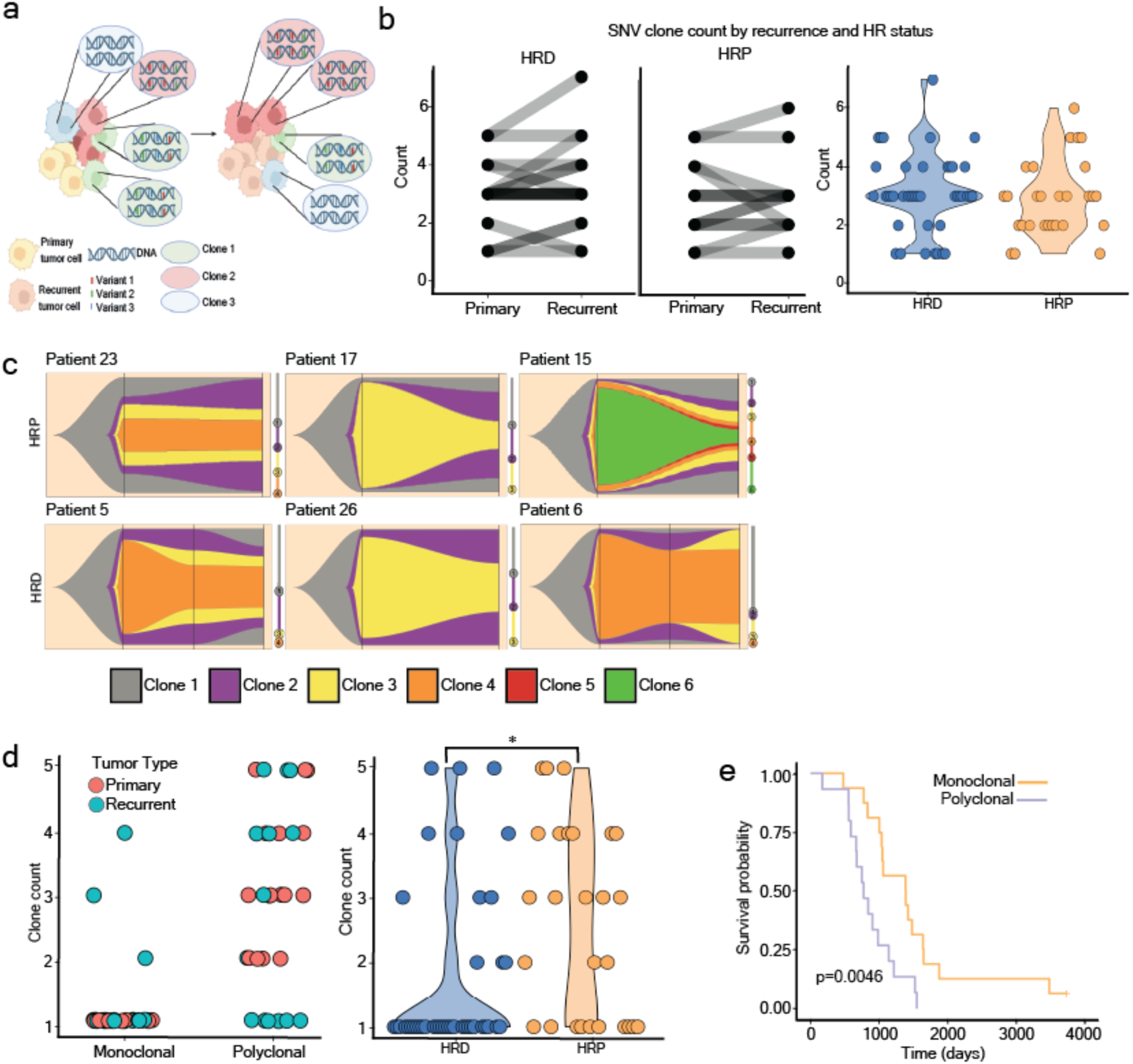
Clonal profiles change little across recurrence. a, Schematic modeling how clonality is measured. Clonal populations will have groups of variants that all increase or decrease in unison as the clone grows or recedes. Measuring the mutation dynamics therefore is a measure of clone dynamics. b, (left) SNV based clone counts, split by HR status. Primary and recurrent tumors are connected by transparent lines. (right) Clone counts per tumor, split by HR status. c, Representative fishplots depicting clonal abundances at indicated times of sampling. Where present, vertical black lines denote a sample collection. Phylogenic trees showing clonal evolution are to the right of each plot. Relative distances between clone nodes on the phylogenic tree indicate the quantity of SNV that separate each clone. d, (left) Primary and recurrent clone counts based on SV dynamics. Tumors are stratified by whether the primary tumor was monoclonal or polyclonal. (right) SV based clone counts, split by HR status. e, Kaplan-Meier plot depicting overall survival of patients with monoclonal and polyclonal primary tumors. *p < 0.05, NS not significant, students T test.

A subset of tumors (13/32) displayed regions of extremely high allele-specific copy number (>5), preventing the estimation of clonality based on all SSMs within these tumors. An average of 4322 SSMs (∼35% of the total SSMs in each tumor) were excluded from clonality estimates in each of these 13 tumors (Supplementary Table 2). These tumors display extreme heterogeneity in the copy number affecting each variant, which is likely a function of increased clonal dynamics and complexity, akin to the adaptive evolutionary state described in^24^.

Using SV data to generate clonality revealed the same trends as SNV based dynamics; few tumors display development of additional clones or purifying selection after chemotherapy, remission, and recurrence. Monoclonal primary tumors tend to give rise to monoclonal recurrent tumors (12/15), while polyclonal primary tumors tend to give rise to polyclonal recurrent tumors (7/13) (Fig 3D). SV breakpoint cancer cell fractions (CCFs) ranged from 0.09-1 and did not correlate with any given type of SV (Supplementary Fig 6B), suggesting little bias was introduced in the cluster determination. Although SVs were highly conserved in some patients (i.e. patient 20 who shared 72% of SV) this was not indicative of low clone count, but rather the conservation of clones (4 clones in primary vs 5 in recurrent). Despite a high correlation between the percent of SNV and SV shared between a patients’ primary and recurrent tumors (R=0.61, p=2.3×10^−11^), there was weak correlation between clonal composition defined by SNV and SV, mostly driven by tumors with one clone (R=0.31, p=0.012; Supplementary Fig 6C, D). Tumors with one identified SNV clone also had one SV clone in 9/10 tumors, while tumors with two SNV clones ranged from 1-5 SV clones (n=12). Tumors with three SNV clones showed the same range (n=28), tumors with four identified SNV clones showed the same range (n=10), and tumors with highly polyclonal SNV (5-8) showed the same range (n=10). Unlike with SNV based analysis, HRD status did affect clonal composition as determined by SV. HRP tumors are more clonally diverse based on SV data, as HRP tumors were more likely to be polyclonal (17/29 polyclonal tumors are HRP) while HRD tumors tend to be monoclonal (29/41 monoclonal tumors are HRD) (Supplementary Table 10).

Polyclonal tumors that contain subclonal (<0.9 VAF or present in a non-truncal cluster) SNVs or SVs tend to exhibit a very high percentage of subclonal variants. A higher percentage of variants are classified as subclonal in HRP than HRD tumors (28.6% SV, 48.7% SNV vs 10.5% SV, 30.3% SNV in HRD), which was expected, as HRP tumors tend to be polyclonal (Supplementary Fig 6C, D, Supplementary Table 10, SV p=3.25×10^−3^, SNV p=0.0434). Tumors showed a very strong correlation between clone count and subclonal SV percentage, which likely drove the high subclonal percentages seen in HRP tumors (Supplementary Fig 6C, E). In our cohort, we noted both an increase in total clones and subclonal variant count in the HRP tumors (Fig 3D, Supplementary Fig 6C, D). Patients with HRP tumors tend to have a shorter progression-free survival in the clinic^25^, suggesting the tumors are more proliferative.

A greater count of SVs in HRD tumors did not correlate with an increased quantity of unique clones (Supplementary Fig 6A, Fig 3D). HRP tumors tended to have a greater quantity of clones (average 2.68 HRP vs 1.73 HRD, p=9.75×10^−3^), despite a smaller quantity of SVs (average SV count; HRP=175, HRD=279, Fig 3B). It is likely that clones in the HRD tumors carry passenger SV introduced in the process of clonal evolution, which contribute to the abundance of total SV. These passenger SV likely increased in abundance during tumor development and recurrence. Patients with polyclonal primary tumors had a significantly shorter progression-free survival than patients with monoclonal tumors, irrespective of HR status (Fig 3E). This trend was more apparent in the HRP tumors, where polyclonal tumors led to a substantially shorter overall survival, which may be influenced by treatment options (Supplementary Fig 7).

### SV signatures reveal three groups of tumors

Five *de novo* SV signatures were identified within our dataset (Fig 4A, Supplementary Fig 8). Signatures 1 and 2 were defined by small and large duplications, respectively. Signatures 3 and 4 were defined by clustered (which may also be classed as copy neutral insertions) and unclustered translocations, respectively. Signature 5 was defined by small deletions (Supplementary Fig 8). Each tumor presented a mixture of SV signatures, so an unbiased hierarchical bayesian clustering method was used to cluster the tumors. We observed closer intra-patient tumor clustering than inter-patient, as previously reported in the cohort using other genomic data types^26^ (p=3.22×10^−14^, Supplementary Fig 9A). As seen in previous work^27,24^, three distinct groups emerged from the resulting dendrogram, tumors defined by signatures 1 and 2 (referred to as duplicators), tumors defined by signatures 3 and 4 (referred to as translocators), and tumors defined by signature 5 (referred to as deletors) (Fig 4A, Supplementary Table 11).

**Fig. 4:**
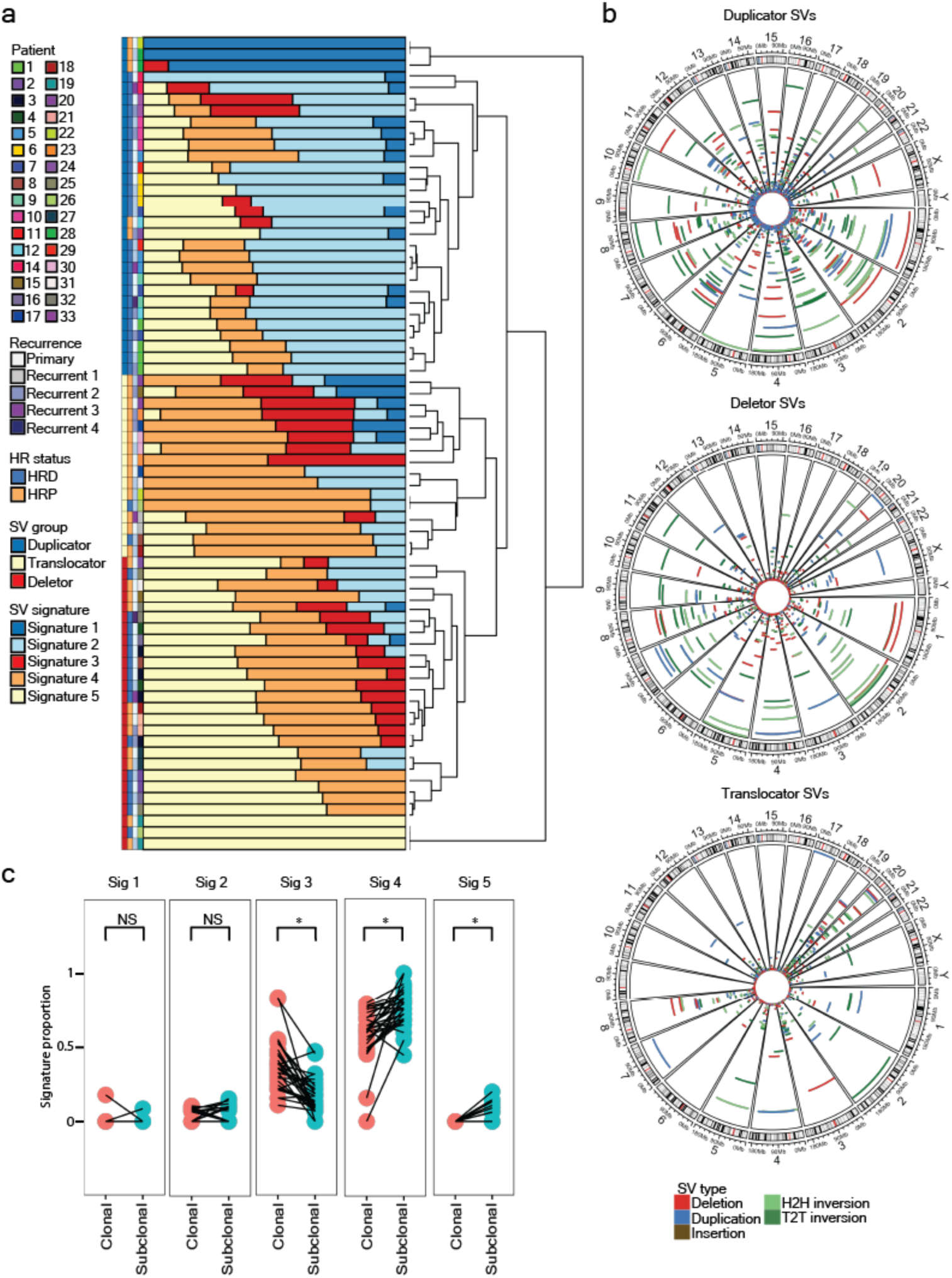
Structural variant signatures identify three distinct SV mutational mechanisms. a, Five *de novo* structural variant signatures applied to each tumor in the cohort and clustered by signature composition. b, Genomic distribution of all SV stratified by SV signature group. c, SV signature proportions in clonal or subclonal populations. *p < 0.05, NS not significant, students T test.

All groups showed significant differences in SV counts with duplicators containing the highest counts, followed by deletors, and translocators (mean 539, 401, 167, p=0.062 del vs dup; p=4.23×10^−7^ del vs trans, p=1.37×10^−4^ dup vs trans; Supplementary Fig 9B, C). As expected, the deletor group contained the highest percentage of deletions, significantly more than any other group (1.67×10^−7^, 8.24×10^−11^, duplicator and translocator respectively). Duplicators contained the highest percentage of duplications (p=1.95×10^−8^, 3.06×10^−5^, deletor and translocator respectively; Supplementary Table 12). Translocators contained the highest percentage of foldback inversions (FBI) and translocations (FBI p=0.018, 0.016, TRA p=0.007, 0.046, deletor and duplicator respectively, Supplementary Fig 9B). These groups were largely, but not completely, defined by germline deleterious variants in *BRCA1* (duplicators, 16/26 with *BRCA1* muts), *BRCA2* (deletors 9/23 *BRCA2* mut), and *BRCA1/2* non-carriers (translocators; 13/18 non-carrier). Only three tumors from carriers of deleterious variants in *BRCA1/2* did not cluster with their group. Consistent with previous reports, the translocators, largely defined by a lack of *BRCA1/2* deleterious variants, had the highest abundance of FBI and lowest patient survivorship (Supplementary Fig 9D)^28^. Patient survivorship was highest in the deletor and duplicator groups, in line with previous reports of carriers of *BRCA1* and *BRCA2* pathogenic variants having increased survival^29^ (Supplementary Fig 9D).

We noted a correlation between the SV signature of a tumor and clonality status. While deletor patients had an even dispersion of monoclonal and polyclonal primary tumors (7 and 5 respectively), duplicators tended to be monoclonal (6/9), and translocators tended to be polyclonal (5/7). Carriers of polyclonal primary tumors had a shorter progression-free survival overall, but this did not consistently apply to the SV groupings. Although the number of cases was low, the progression-free survival in the duplicator and translocator groups was not affected by the presence of multiple clones. The deletor group exhibited drastically shorter overall survival intervals if a patient had a polyclonal tumor (p<0.0001, Supplementary Fig 9E). Additionally, the shorter overall survival seen in polyclonal tumors was primarily driven by the deletor group, in which the overall survival was more than double the length for patients presenting monoclonal tumors, as defined by SV, versus polyclonal (∼750 vs ∼1500 days Supplementary Fig 7, Supplementary Fig 9E, p=7.9×10^−4^).

The distribution of SVs across the genome was unique to each SV signature group (Fig 4B, Supplementary Table 13). As the translocator group was heavily composed of HRP tumors, the SV distribution closely matched the HRP SV profile, while the deletors and duplicators bore aspects of the HRD profile. Translocators lack large SV in chr 9 and 10, while having an abundance in chr 19 and 20. Deletors display the most abundant large SV in chr 9 while lacking almost all large SV in chr 14, 17, 18, and 19. Duplicators display all the chr 17 q arm inversions seen in the primary/HRD group while lacking SV in chr 19 and 20 (Fig 4B, Supplementary Fig 2B, D).

SV signatures were largely conserved across recurrences. In only four cases did a primary tumor cluster in a different group than the patients paired recurrent tumor, and in 3/4 cases the group switching was between the two groups that were polyclonal (deletors and translocators). The fourth case, patient 24, transitioned from duplicator to deletor. Interestingly, this tumor pair showed an expansion of clones which is consistent with the expansion of SV and SV signature alteration seen in this patient (Supplementary Table 10, Supplementary Table 11). None of the patients with tumors in multiple groups had altered HR status, and both tumors from the patient with altered HR status (patient 31) belonged to the deletor group. In line with clonality, this lack of group alteration seems to suggest that the tumor’s mutational profile is established early in tumor development and conserved throughout treatment.

### SV signatures and clonality bias

The five identified *de novo* SV signature proportions were measured in each polyclonal tumor (n=29) using subclonal or clonal SVs separately (12 HRD, 17 HRP). Signature 3 (defined by clustered translocations) was identified in all 29 tumors, and was significantly higher when clonal SVs were used (p=2.2×10^−6^). Signature 4 (defined by unclustered translocations) was also identified in all 29 tumors but was a significantly higher proportion in each tumor when subclonal SVs were used (p=1.95×10^−5^). The enrichment of this signature may be due to increased accuracy in identifying translocation breakpoints, allowing these variants to pass the stricter filtering needed to identify clonality. Signature 5 (defined by small deletions) was identified in 8 tumors and was only present when subclonal SVs were used (p=6.06×10^−3^). No significant differences in signature proportions were identified for the remaining signatures (Fig 4C, Supplementary Fig 10).

### Few genes are commonly affected by SV

While SV sizes were relatively similar within the deletor and duplicator groups, the distribution of SVs in the genome were distinct. Small SV were found throughout the genome in each group, but large SVs displayed an accumulation SV in chr 17 in the duplicator SV group (4.49% of SV within the duplicator group), which was not seen in either deletors (1.2%) or translocators (0.46%). Most of the chr17 variants in the were found in the q arm, which was consistent across SV groups, with an average of 77% of chr 17 SVs being found on the q arm. Translocators displayed accumulation of large and very large SV on chromosomes 19 (31.03% of large/very large SV) and 20 (12.1%), which were relatively unaffected by large and very large SVs in the deletor and duplicator groups (Supplementary Fig 11). Deletors did not show a notable accumulation in any chromosome, relative to the other groups. Chromosomes 1, 2, 3, 5, 7, and 8 were the few chromosomes that possessed at least 5% of large SVs across all SV signature groups (Fig 4B).

Genes commonly affected by SVs shorter than 5 MB were rarely observed. The most commonly disrupted gene was *CNTNAP2*, which was deleted in 20/72 tumors. This may be a function of its size, as *CNTNAP2* is approximately 2.3 Mb in length. Although individual genes were infrequently altered by SVs, pathway analysis identified genes from cellular metabolism or cell cycling and tumor suppressor genes to be enriched for SVs (Supplementary Table 14).

### Long read sequencing validates most short read calls while elucidating complex rearrangements

We performed ultra-long ONT sequencing in five tumors for which sufficient tissue was available to perform high molecular weight DNA extraction. ONT sequencing yielded an average output of 80.06 gigabases, with an average N50 of 55.82 kb per tumor sample. Using phased reads, an average of 1696 (median 615) SVs were called in each sample with a median phase block N50 of 11 Mb. An average of 32 SV clusters occurred per sample, as determined by Severus^31^ (Supplementary Table 15, 16). Fig 5A depicts one such cluster in the recurrent tumor of Patient 14, where chromosomes 3, 8, 17, and X display multiple breakpoints generating new derivative chromosomes. The phased long read sequencing allowed us to identify the haplotypes that in which each breakpoint was present and identify these new derivative chromosomes arising from complex structural variants. Seven of the 14 breakpoints in the cluster lie in long interspersed nuclear elements (LINES) while both of the chromosome 3 to chromosome 8 translocation breakpoints lie in short interspersed nuclear elements (SINES). When regions ± 200bp around each breakpoint were analyzed, we found that each breakpoint pair shared at least one region of 10 bp in length with exact homology (Supplementary Fig 12), giving some insight into the mechanism of donor site selection.

**Fig. 5:**
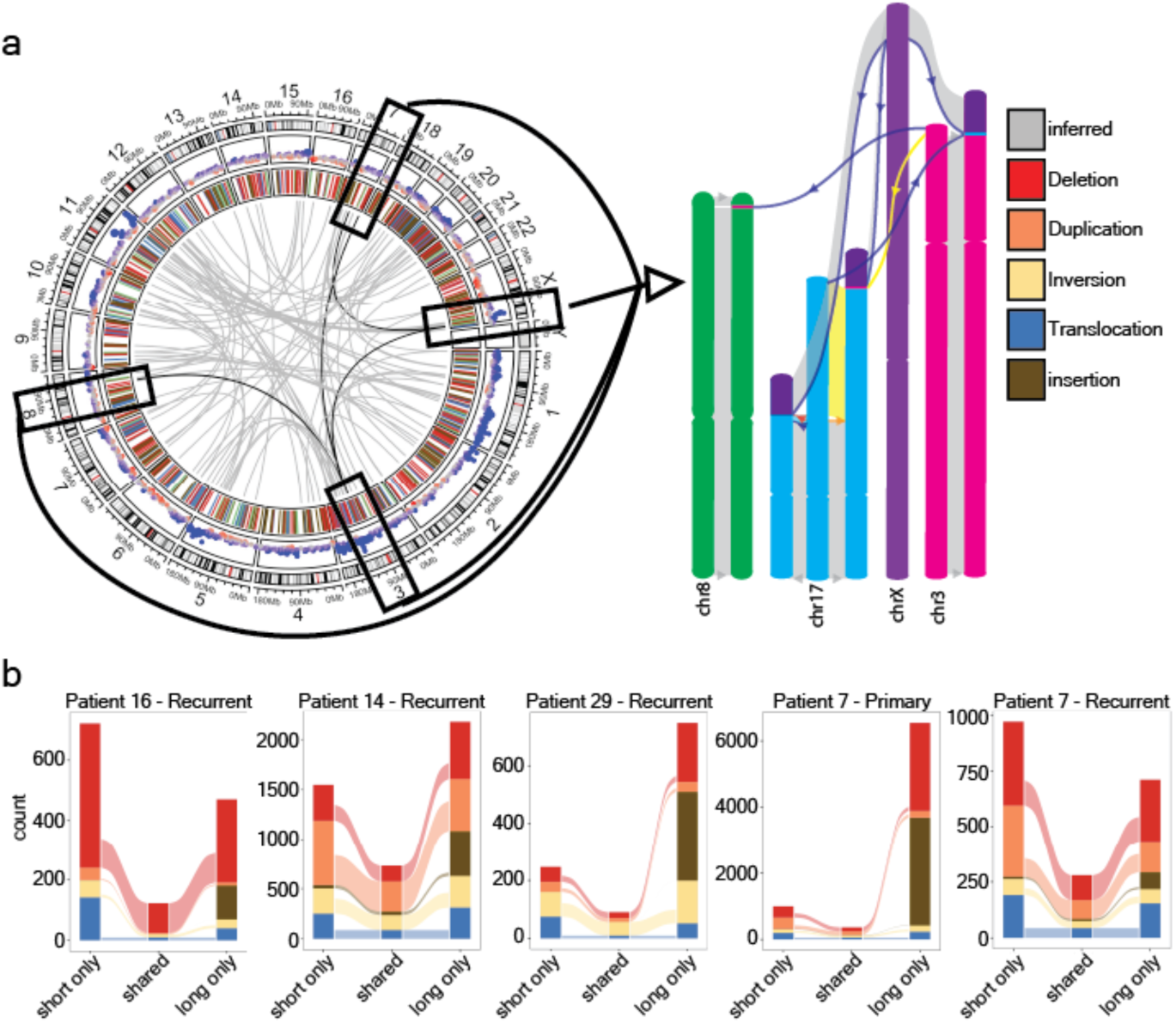
Haplotagged long reads identify complex chromosomal rearrangements. a, SVs identified with long read sequencing from the recurrent tumor from patient 14. (Left) Circos plot showing copy number and structural variants generated from short read sequencing for a representative SV cluster. (Right) Diagram depicting derivative chromosomes generated as a result of a single SV cluster, as they are observed in the tumor using phased long read sequencing data. Colored arrows depict the type of SV and the donor chromosome. b, Alluvial plot showing the quantity of variants shared amongst short and long read sequencing.

There were significant differences identified in the number of SVs in short read and long read sequencing. On average 56% of SVs identified by short read sequencing were also identified by long reads, which increases to 63% if translocations are excluded (Fig 5B, Supplementary Table 17, Supplementary Fig 13A, B). Short reads identified many more duplications >5Mb, whereas long reads identified far more insertions (Supplementary Fig 13C). As expected, we identified increasing counts of SVs as we acquired more reads from each tumor, highlighting the abundance of subclonal rearrangements that are only present on a small number of reads (Supplementary Table 16). Although most SVs were replicated, long read sequencing identified several additional tumor suppressor gene mutations missed by short read sequencing. In patient 29 we identified a 133 Mb deletion covering *VHL* (19% VAF), in patient 7’s primary tumor a 351 bp deletion in *NF1* (24% VAF) and 335 bp insertion in *RB1* intron (20% VAF), and in patient 16’s recurrent tumor a 41Mb deletion covering *RB1* (25% VAF) (Supplementary Table 17).

Known *BRCA1/2* deleterious variants and SVs detected on chromosomes 8 and X using short read sequencing were investigated further in the long read sequencing data. Patient 7 is a heterozygous carrier of a *BRCA1* deleterious variant (Supplementary Table 1). While short read sequencing did not suggest a reversion, long read phased data showed reads mapped to the deletion haplotype that bore *BRCA1* reversion, suggesting clones with a reversion may have arisen (Supplementary Fig 13D). Chromosome 8q amplification, which harbors the MYC locus, was one of the few copy number profiles shared across many of the tumors in our cohort (Fig 2A). Fourteen 1 kb-1 Mb duplications with ≥ 2 fold changes in relative depth were verified at this locus (Supplementary Fig 13E), twelve of which were also detected in short read sequencing and shared exact breakpoints with the long read data. However, 4/12 shared duplications were identified as templated insertions and 6 of the 12 copy number gains were tandem duplications (Supplementary Table 18). None of these SV fully explained the increased copy numbers observed in the MYC locus. The progressive copy number gains without accompanying SVs suggests we need greater depth to capture the full range of genomic instability at this locus, or that the increased depth is the result of ploidy alterations. Chromosome X displayed frequent large scale copy number losses based on short read data, as shown in Figure 2A. Two tumors used for long read sequencing displayed copy number <2 across the X chromosome with short read sequencing. Both tumors were profiled at ≥ 20X depth with ONT long reads, and had sufficient coverage for phasing analysis^32^. Neither tumor was found to have loss of heterozygosity on the X chromosome, rather SVs were interspersed across the chromosome and affected both haplotypes. Interestingly, both tumors were found to have “haplobiased” X chromosomes^32^, with a majority of haplotype 1 SVs occurring in the first 60Mb, few SVs found in the 60-100 Mb region, and a majority of haplotype2 SVs occurring the 100-155 Mb regions (Supplementary Table 18).

## Discussion

Patients diagnosed with HGSOC are presented with limited treatment options and a majority relapse with progressive tumors. Deeper understanding of the molecular signatures that underlie mutational changes that contribute to genomic instability is imperative to improve patient outcomes. Biopsies of recurrent tumors are rarely collected^9^, making this cohort a highly valuable and novel resource. Previous reports have focused on utilizing ascites as a proxy for recurrent HGSOC tumors, and have relied on SSM analysis to investigate origin and resistance mechanisms^11,19^. The low rate of *TP53* mutations detected in ascites samples serves as a molecular indicator of their unsuitability to represent solid recurrent HGSOC^33^. Thus, understanding the role of SVs in primary tumors and matched recurrent tumors has not been investigated due to the limited availability of paired chemo-naive primary and recurrent tumors collected at time of relapse. In the present study, we examined a cohort of 32 HGSOC patients with paired chemo-naïve and chemoresistant tumors, as well as germline DNA. Our analyses are consistent with previous reports where few genes, outside of *TP53*, are frequently mutated across HGSOC tumors. However, likely due to the increased sequencing depth used in our study we identified a novel commonly mutated gene, *MDC1*, that may confer homologous recombination repair deficiency outside of the standard *BRCA1/2* paradigm by inhibiting the formation of the *BRCA1/2* complex.

*MDC1* serves as a docking platform to promote the localization of various DNA damage response components, including *BRCA1*, to DNA double-strand break (DSB) sites^34^. Assembly of the MDC1-histone (γ-H2AX) complex seems integral to retention of most other proteins in these regions and loss of *MDC1* impairs DSB repair^35^. Importantly, mutations in *MDC1* were almost exclusively located in the PST repeat domain region (Supplementary Fig 2), which specifically binds to the DNA-dependent protein kinase (DNA-PK) and regulates autophosphorylation of DNA-PK^36^. Furthermore, mutations in the PST domain have been observed in melanoma, bladder cancer, prostate cancer, and non-small cell lung cancer patient samples^34^. Taken together, the specific location of the *MDC1* mutations in tumors in this study suggest impaired error-free DSB repair due to impaired DNA-PK binding ability and error-free DSB repair.

Although functional validation of these point mutations was inconclusive, the tendency for these mutations to be in HRP tumors with pronounced genomic instability cannot be understated. Eight of the nine tumors with identified *MDC1* mutations had no germline deleterious variants in *BRCA1/2*, which may explain how these tumors became genomically unstable. As HR deficiency is associated with chromothripsis^37^, the higher incidence of chromothripsis observed in *MDC1* mutants may be driven by *MDC1* mediated HR deficiency. Similar to *BRCA1/2* loss, the inactivation of *MDC1* induces sensitivity to DNA damage^38,39^, which could explain why we observed a higher proportion of *MDC1* loss in primary tumors (Supplementary Table 3). Outside of *MDC1*, homologous recombination was stable throughout disease. Only two patients had an altered disease status (1 reversion, 1 in-frame deletion) across the course of disease. Although the reversion rate is estimated to be up to 26% in HGSOC ^40^, we did not observe this in our cohort, possibly due to sample size.

Similar to Gull et al 2022^15^, somatic mutation profiles were largely conserved throughout the disease course from primary to recurrent tumors. SSM and SV burden were not significantly different in tumors from different time points in disease course from the same patient. Interestingly, large SVs affecting chromosomes 19 and 20 in HRP tumors were not unique to either primary or recurrent tumors. This implies these variants may be important for both tumorigenesis and maintenance. Large SVs on chromosome 17 were identified in almost 40% of HRD tumors (Fig 2E). These SV are predominantly inversions, making it difficult to infer their functional effect, but suggesting that altering expression of genes in this region may aid in tumor and clonal expansion, but an outright loss is not beneficial. Inversions have also been shown to be associated with DNA methylation changes^6,16^ suggesting carriers of these inversions may have unique responses to environmental changes or therapeutic intervention. The proportion of shared events across HRD tumors was lower than HRP tumors, suggesting that the mutational landscape is relatively conserved across variant types, likely due to a preservation of clonal dynamics.

An assumption of a single initiating cancer clone that gives rise to a diverse polyclonal tumor that is subjected to purifying selection is the standard dogma for many cancer types^41,42^. However, this mechanism was not observed in this study, wherein we observed sustained clonal and subclonal populations throughout chemotherapy, surgical resection, and tumor regrowth. This suggests that the drivers for chemoresistance are present within the primary tumor and chemotherapy does not drive the development of new clones, but rather filters out the clones that were chemo-sensitive. We did observe that subclonal SV tend to be deletions while clonal SV tend to be duplications (Fig 4C). Duplications are less likely to lead to cell death^43^ while deletions, particularly in DNA repair genes, lead to genomic instability^44–46^. These deletions are believed to be random and reflect the evolutionary process of increasing fitness.

Additionally, high grade serous ovarian tumors are frequently polyclonal, and their persistence suggests tumor clones act synergistically to facilitate tumor growth. Previous studies with smaller cohorts have seen polyclonal seeding of metastatic sites^7,24^ which was also observed in our cohort. Patients with polyclonal primary tumors (as defined by SV) tended to have polyclonal recurrent tumors with similar clonal compositions, and also displayed a shorter progression-free survival than patients initially presenting with monoclonal tumors (Fig 3E). A more diverse set of clones would set up the tumor to handle chemotherapeutic insults better, which may partially explain the higher mortality seen in HRP tumor bearing patients^47^. Consistent with our cohort, high levels of intratumoral heterogeneity (i.e. increased clone counts) have been shown to diminish immune responses and survival intervals^48^.

Structural variant signatures, largely re-capitulating the genomic impacts of the presence of loss of *BRCA1/2* function, were also associated with reduced survival (Fig 4A, D). Previous publications have identified the relationship between *BRCA1* loss leading to increased duplications, and *BRCA2* loss leading to increased deletions and their phenotypic outcomes^49^. However, some *BRCA1/2* non-carrier patients share similar SV and SV signatures to these two groups, and potentially have similar molecular profiles. *BRCA1/2* non-carrier patients that display deletor/duplicator signatures may benefit from the same interventions as *BRCA1* and *BRCA2* deleterious variant carriers.

Recent reports have indicated that short reads are capable of capturing a vast majority of the complexity reported in long reads, and that the main advantage of long read sequencing is phasing^50^. It was unclear if these results would be replicated in genomically unstable HGSOC tumors. Short read sequencing successfully identified many of the large SVs that accompanied copy number changes, but was less accurate at calling >300 bp deletions that had a less pronounced copy number effect. Insertions, templated and non-templated, are particularly difficult for short reads to identify, which led to overcalling duplications based on local sequencing depth from short read data (Supplementary Fig 13B, C). As expected, most short (>100bp) deletions are accurately captured. Around half of the deletions and duplications (50%, 46% respectively) were confirmed by long reads, suggesting that the structural variant signature groups dominated by deletions and duplications likely represent true genomic processes. Interestingly, the most replicated >5MB SV type was inversions, which are not typically associated with copy number changes, and therefore have fewer lines of evidence in short read sequencing (Supplementary Fig 13B). While just over half of the short read variants were validated by long read sequencing, it is important to note that these data were generated from adjacent tissue punches, collected at different times, with different extraction methods. Given the high level of conservation of somatic variants across time points in the disease course, even after many treatments with chemotherapy, it seems unlikely that sampling variation at a single time point would significantly contribute to the large proportion of variants uniquely identified by long read sequencing, although it cannot be entirely ruled out. It is also feasible that multiple short variants on separate haplotypes were incorrectly called as a single long variant in short read data, inflating the number of unique variants identified with long read sequencing data.

While our study does represent one of the larger cohorts of matched primary and recurrent tumors with genomic profiling, we are still underpowered to comment on the effect that individual chemotherapy regimens may have had on the molecular profile of tumors. As patients’ disease progressed additional lines of treatment were added, making the treatment regimen beyond the first recurrence extremely heterogenous across the cohort. To separate out the variants that drive recurrence and those associated with individual chemotherapy agents, we will need a larger cohort. Still, we have shown that inter-tumor heterogeneity is pronounced in HGSOC and that tumors from the same patient more closely resemble one another than any other, highlighting a scenario by which each patient develops their own largely private set of somatic mutations, albeit through shared mutational mechanisms. This underscores the need for individualized patient care based on tumor profiling to identify alternate therapies that may target specific molecular vulnerabilities in each patient.

## Methods

### Cohort description

Fresh-frozen matched primary (n=34) and recurrent (n=38) high-grade serous ovarian cancer specimens and DNA from thirty-three women diagnosed with high grade serous adenocarcinoma were included. Specimens were all identified in the Cedars-Sinai Medical Center Women’s Cancer Program Biorepository (IRB #0901). All patients gave informed consent to participate in the study. 11 participants were known carriers of deleterious *BRCA1* and/or *BRCA2* germline variants, the remaining participants were not carriers of deleterious variants in any known high or moderate effect breast or ovarian cancer risk genes. All patients were diagnosed with stage III or stage IV disease and underwent primary optimal surgical cytoreduction (to less than 1 cm residual disease) prior to administration of combination chemotherapy with platinum and taxane between the years of 1990 and 2014. For each patient, detailed clinical data were available including clinical genetic testing results, dates of original diagnosis and each subsequent recurrence, treatments administered throughout their disease course, operative and pathology reports, and other clinicopathologic variables including other cancer diagnoses and comorbidities. These details can be found in Supplementary Table 1. The tumor samples were sequenced at a depth of 80x (mean 67x; Supplementary Table 2). Stranded total RNA sequencing (RNA-Seq) was performed and generated a mean of 335 million reads per library, and whole genome bisulfite sequencing (WGBS) was performed to capture methylation at a mean depth of 30x, as previously reported^15^.

### Specimen acquisition and preparation

Germline DNA was extracted from whole blood drawn into 4mL EDTA treated collection tubes at the time of debulking surgery. DNA was extracted with the Qiagen DNEasy Blood & Tissue Kit (Qiagen, MD, USA) and quantitated with the Quant-IT dsDNA Broad Range kit on a QuBit (Thermo Fisher Sci, CA, USA). Fresh-frozen tumors were embedded in optimal cutting temperature (OCT) compound, bisected and mounted and two slides were made for hematoxylin and eosin (H&E) staining. All slides were reviewed by a single pathologist to identify regions enriched for epithelial carcinoma (avoiding regions enriched for stroma). These regions were sampled in a single core of approximately 50 mg collected on dry ice. Each core was divided into three pieces, one of which was used for genomic DNA (gDNA) extraction using the Macherey-Nagel Nucleospin DNA Kit (Machery Nagel, Germany). Remaining tissue was used for gene expression profiling and proteomics profiling (described in Gull et al, 2022 and Dabke et al, *Under review*). DNA samples were quantified using the QuBit (Thermo Fisher Sci, CA, USA) to measure the content of double stranded DNA and quality was confirmed using the Agilent TapeStation (Agilent, CA, USA).

### Library preparation

gDNA was diluted in water to a concentration of 50ng/ul and a total of 1ug of DNA was shipped to Fulgent Therapeutics (Temple, CA, USA) for whole genome sequencing. DNA samples were sheared using the Covaris m220 to fragment size 400-500bp, which was confirmed with the Agilent TapeStation. Agencourt AMPure Beads were used for fragmented DNA purification and size selection. The NEBNext® Ultra™ II DNA Library Prep kit | NEB and Illumina TruSeq Index Adapters were used to generate PCR-free sequencing libraries following manufacturers protocols.

### Whole Genome Sequencing

Libraries were sequenced on either a single lane (germline DNA) or two lanes (tumor DNA) of the HiSeqX (Illumina, CA, USA). Fastqc and Picard Tools (*CollectAlignmentSummaryMetrics*, *CollectWGSMetrics*, *CollectGcBiasMetrics*) were used to generate QC reports that were visualized in MultiQC. Samples with insufficient read depth were excluded and an additional sample aliquot was repeated using the methods described above. Sequencing reads were aligned to hg38 using a custom BWA-GATK pipeline in the Cancer Genomics Cloud, in which indels were realigned with IndelRealigner, base quality estimated, and base quality recalibrated with BaseRecalibrator^51^. Coverage metrics, GC bias, and mismatch rate and chimera rates were collected with picardtools^52^, coverage was calculated using bedtools genomecov^53^. These metrics are summarized in Supplementary Table 2. Matching germline and somatic samples were verified using PLINK^54^.

### Mutation Calling

A consensus pipeline for calling indels and SNV was constructed using Mutect2^51^, Strelka2^56^, VarScan2^57^, and Somatic Sniper^58^, in which a variant must appear in at least two separate callers to be retained. Variants with fewer than 5 supporting reads were removed. Further filtering was applied to SNV wherein a base quality score of lower than 20, those located in regions of the lowest 1% mappability score^59^, and variants within intervals included in the Encode Blacklist^60^ were removed. Samples that did not have a somatic driver mutation identified in the automated variant calling and consensus pipeline in *TP53* were manually inspected in IGV^61^ according to best practices^62^.

### Short read SVs: GRIDSS

The GRIDSS^63^ suite was utilized to generate SV and copy number calls genome-wide. This analysis suite includes several analysis tools, including Gripss (a filtered version of the GRIDSS SV calls), Cobalt (which determines read depths), Amber (which determines BAF), Purple (PURity and PLoidy Estimator), and Linx (grouping and clustering SV events)^63^. A workflow chaining these individual programs was designed within the Cancer Genomics Cloud that incorporated Gripss v 2.10.2, Cobalt v 1.1, Amber 1.9, Purple v 2.51, and Linx v 1.14. Tools were run according to standard settings.

### PCGR: Mutation rate and chromothripsis

We utilized the dockerized version of PCGR (version 0.9.2)^64^ to estimate the mutation rate and MSI status of each tumor sample in our cohort. Chromothripsis was determined via the Shatterseek program^65^, in which regions of at least three oscillating copy number profiles are colocalized with at least four nested structural variants are marked as likely chromothriptic. The output of Shatterseek was manually curated for accuracy of copy number oscillation and SV events, with only those meeting the copy number and SV criteria being retained.

### Significantly mutated genes

Genes mutated at a significant rate above the background mutation rate were identified using the ActiveDriverWGS tool^66^. A list of all somatic variants was cross referenced with a table of all coding regions from the Gencode v34 (gencode.v34.annotation.gtf), which was downloaded from the UCSC table browser. Mutated genes appearing with at least eight observed mutations and a p value <0.005, and an FDR rate lower than 0.005, were determined to be significantly mutated. Mutations were then manually inspected to verify the surrounding regions were not subject to misalignment or high discrepancy regions^62^.

### Amplicon sequencing

Somatic mutations in *MDC1, TP53, KRAS, BRCA1, BRCA2, RB1, MYC*, and *CCNE1* were sequenced using targeted amplicon sequencing. Tumor derived DNA used for short read Illumina sequencing was aliquoted (50ul of DNA at 18ng/ul) and mixed with 10ul of 2x Ampliseq custom primer pool (REF:20020495, lot:10841836, Illumina, CA, USA), and 5ul of Ampliseq HiFi mix were added per sample. The Ampliseq custom primer pools contained 100ng of each primer set (Pool 1: 347 primer pairs, Pool 2: 342 primer pairs). Libraries were quantified and pooled and sequenced on the Illumina NovaSeqX+ to an average of 5M reads and 9562x coverage. Sequencing reads were aligned with BWA^92^ and variants called in tumor only mode using Mutect2^51^.

### PRS

The polygenic risk score (PRS) of an individual *i* was calculated by a linear function of 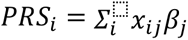, where *x_ij_* represents the number of effect alleles at the *j*th SNP for the *i*th individual, and *β_j_* represents the log odds ratio of the *j*th SNP. Genotypes are denoted as *x*, with values 0, 1, or 2, indicating dosage of the effect allele. The PRS model for overall EOC comprises 15 SNPs, which were obtained from^67^, along with their corresponding log odds ratio. To calculate PRS for each patient, we used PLINK 2.0^68^.

### PIK3CA/B amplification

Copy number profiles for genes were generated by utilizing b allele frequencies analyzed through the Purple function in the GRIDSS suite^63^. If a gene contained a copy number that rounded up to 6 or higher it was deemed an “amplification”.

### Whole genome duplication

Copy number profiles generated by PURPL (within the GRIDSS suite) were used to determine if whole genome duplication (WGD) had occurred. WGD was defined as >70% of the genome displaying a copy number profile above 4.

### Shared SNV, indel, and SVs

SNV and indel calls generated by the consensus pipeline were intersected with bedtools^53^ intersect function with exact matching to determine quantities of private and shared variants. SVs were intersected with a minimum overlap of 70% between both SV for them to be counted as the same variant. Translocations with both breakpoints within 10bp of another translocations breakpoints were merged and considered as a single event.

### Homologous recombination status prediction

HRD was determined by considering several lines of genomic evidence; germline carrier status of deleterious variants in *BRCA1/*2, somatic promoter hypermethylation of *BRCA1/2*, *RAD51C,* or *RAD51D*. CHORD score was determined using the Classifier of HOmologous Recombination Deficiency (CHORD) score^17^. SNV and INDEL information extracted from Mutect2 VCF files, SV information extracted from gripss VCF files, the filtered high-confidence output from GRIDSS, were used as inputs for the tool. Default parameter settings were used for the analysis and resulting classification labels.

### Clonality analysis

To determine allele and clone specific copy number alterations along with clone proportions we utilized HATCHet^69^. This gave clonal composition and proportions for each mutation present within the sequencing data. The clonal matrix was cross referenced with the consensus file to remove the mutations that were more likely to be false positives. DeCifER^70^ determined the clone that each variant belonged to and the descendant cell fraction (DCF) of each variant. Per DeCiFer’s documentation, only mutations that appeared in both the primary and recurrent tumor were considered for clonal evolution. Unique state trees were generated for each tumor sample where possible. ClonEvol^71^ interpreted the clonal structure, as defined by DeCifER, and the mean DCF of each clone, to generate clonal evolutionary structure. Fishplots for each tumor were generated by providing ClonEvol with DeCifER clonality results. True cluster DCFs were used for SNV estimations, in the cases when no consensus model could be found the point estimate DCF was used to guide fishplot generation. Resolving the clonal structure was not possible for 6/33 samples, which may be due in part to single site sampling. In order to determine how SV abundance and CCF of SV breakpoints affected clonal composition, we used SVclone^72^. Currently SVclone cannot accurately profile the temporal progression of SVs, therefore tumors from the same patient were analyzed separately.

### SV signatures

Structural variant calls generated by GRIDSS were further analyzed by the “Palimpsest” R ^30^ package. The package determined if SVs were clustered by initiating events or occurred in an unclustered manner. Total SV counts and SV sizes were utilized to generate denovo SV signatures. Each tumor was analyzed for signature content and assigned a signature content value for each signature identified in the cohort. Within the data set there were several tumors that only displayed a single signature while others displayed up to four different signatures per tumor. Tumors were clustered with a euclidean clustering algorithm and the resulting dendrograms were used to determine SV signature groups.

### Clonality defined by SVs and subclonal signatures

Structural variant calls generated by GRIDSS, in the VCF format, along with bam files were input in the SVclone^72^ program. Read information present in the bams was used to adjust VAF for each breakpoint. After a filtering step, the SVs were clustered according to CCF, copy number intervals, and tumor purity. The Ccube clustering method was applied to the CCF adjusted breakpoints to determine cluster assignment for each SV. A joint SV and SNV clustering method was applied to the sample. SVclone does not currently support multi-tumor analysis, and other standard tools, such as pyclone^74^ or DeCifER^70^, do not officially support SV clonal analysis. Therefore, we did not analyze temporal evolution of clones as defined by their SV profiles. Variants assigned to clone 1 were determined to be clonal while variants below a VAF of 80% or assigned to any other clones were designated as subclonal. We applied the five denovo signatures found in our total SV dataset to both the clonal and subclonal SV datasets. A total of 24 tumors were analyzed for subclonal and clonal SV signatures. Signature abundances were compared between clonal and subclonal signatures, with a p value <0.05 considered significant. SVs were visualized as circos plots generated using the circlize R package^75^. SVs are arranged according to the percent of the chromosome that they affect.

### Pathogenic structural variants

Svanna^76^ was used to annotate SVs with functional annotations to predict deleterious effects on coding regions at SV sites. SVanna generates a ranked list based on intersections with predicted functional domains of coding regions, and membership in the top 10 is highly correlated with pathogenicity of a structural variant. To further select for highly pathogenic SV we retained only the top 5 predicted pathogenic SVs from each tumor. Predicted pathogenic SVs were filtered to include SVs ranked in the top 10 with a Svanna score > 2500. This cutoff represented an inflection point in the distribution of the scores across the annotated SVs in the study.

### Statistical analysis

Statistical tests were performed in R using the rstatix package^55^. All reported statistics were two tailed students T tests, unless otherwise stated. Survivorship analysis was performed using Kaplan-Meier plots that were generated using the “survival” R package^73^.

### GO analysis

GOrilla^77^, was used to annotate gene ontology of coding regions impacted by SVs. To create the input, all structural variants were intersected with the Gencode v34 gene annotations, available through the UCSC genome browser’s^78^ table browser^59^. This produced a data table with genes affected and the type of structural variant affecting them. To keep likely functional effects separate, we separated duplications with breakpoints within the coding region from duplications covering the entire gene. The list of genes affected by SV was sorted by the number of unique occurrences each gene was impacted by a given SV type across each tumor in our cohort.

### Copy number depth profiles

Read depth information was extracted from the bam files using the mosdepth tool^79^. The tool measured depth from BAM files in genomic windows of 10kb. Relative depth was determined by dividing the depth at a given region within the tumor sample by the total depth of the same region within the corresponding normal.

### SNV Count: genomic region cartoon

*MDC1* gene region figure was produced using pyGenomeTracks^80^ using python version 3.7and RefSeq Gene hg38 gtf. Genome was plotted from chromosome 6 position 30695807 to 30720000.

### Long read sequencing

An additional punch was collected from epithelial enriched regions of the same fresh frozen specimen and DNA extracted using the Monarch® HMW DNA Extraction Kit for Tissue (New England Biolabs, MA, USA) and quantified using the Qubit method described above. Samples were profiled for fragment analysis used the FemtoPulse (Agilent, CA, USA). Ultra long sequencing libraries were generated using the Ultra Long read Kit (ULK114) for tumor DNA, and the Ligation Sequencing Kit V14 (SQK-LSK114) for germline samples and sequenced on the ONT Promethion FlowCell vR10.4.1 on the Promethion P24 (Oxford Nanopore Technologies, Oxford, United Kingdom). Germlines were processed on as many flow cells as possible to achieve 100GB of data. Each tumor derived library was sequenced on a single flow cell. Tumor derived libraries with less than 50GB of data generated from a single sequencing run, corresponding to ∼15x coverage, were excluded. Reads were aligned to hg38 using the map-ont option in minimap2^81^. Clair3^82^ identified SNPs in the long read data, which was used in haplotagging the bam file with Longphase^83^. N50 of phase blocks was determined using a script provided in the longphase github. Haplotagged somatic and germline bam files were used by Severus^31^ to identify somatic structural variants and variant clusters. Severus default parameters of at least 3 supporting reads, SVs of ≥50 bp, and a minimum VAF of 0.05 were used. Shared SVs between ultra long and short read sequencing datasets were determined by intersecting SVs called in each dataset and retaining only those variants that had ≥80% overlap between each dataset.

### Protein extraction

A 3 mm core of approximately 50 mg was then extracted from a region of pure epithelial carcinoma from the intact frozen tumor, with care not to include any of the surrounding stroma. These “pure tumor” tissue cores were used for subsequent proteomic analyses. We also extracted regions adjacent to the “pure tumor” tissues which were either a mix of tumor and stromal samples (labeled as 50% stromal content) or stromal samples (labeled as 100% stromal content). Frozen tumors and adjacent stromal samples were thawed on ice, 200–400 μL lysis buffer (6 M Urea + 0.1% Rapigest) was added, and tissues were homogenized with a polytron. Proteins were extracted by high-pressure barocycling on a Pressure BioSciences instrument, model 2320EXT (PBI is Easton, MA) at room temperature, ramping pressure to 45 kPSI, holding for 50 s, returning to atmospheric pressure for 10 s, and repeating for 60 cycles. Protein concentration was assayed using the Pierce BCA assay. Three data sets were prepared, a pool of all available protein lysates (pooled homogenate acquired at 2, 4, 8 μg, each with three technical replicates), a pilot set of tumor samples from 6 HGSOC tumors with either 2 or 3 technical replicates each (*n* = 14) and a clinical data set of tumor and tumor adjacent stromal samples from 16 individual patients, 4 of whom had more than one primary tumor or adjacent stroma sample available for profiling (*n* = 41).

### Proteomics data acquisition

Data acquisition was performed as described previously^87^. Briefly, 4 ml of the digested sample was injected directly into a 200 cm micropillar array column (uPAC, Pharmafluidics) and separated over 120 minutes reversed-phase gradient at 1200 nL/min and 60 C. The eluting peptides were electro-sprayed through a 30 um bore stainless steel emitter (EvoSep) and analyzed on an Orbitrap Lumos using data-independent acquisition (DIA) spanning the 400-1000 m/z range. After analysis of the full m/z range (40 DIA scans), a precursor scan was acquired over the 400-1000 m/z range at 60,000 resolution. To construct a comprehensive peptide ion library for the analysis of human ovarian cancer, we combined several datasets, both internally generated and from external publicly available resources. First, we utilized a publicly available HGSOC proteomics experiment by downloading raw files from the online data repository and searching them through our internal pipeline for data-dependent acquisition MS analysis as described in Parker et al.^84^ and performed by others on the same publicly available dataset ^85^. Database searches for both the internal and downloaded external datasets utilized human protein sequences defined in a FASTA database of Swiss-Prot-reviewed, Human canonical reviewed proteome containing 20,406 protein sequences that were downloaded July 2019 and appended with Biognosys indexed retention time (iRT) peptide sequence (Biognosys, Schlieren, Switzerland) and randomized decoy sequences appended. A final, combined consensus spectral library containing all peptide identifications made between the internal and external datasets was compiled. Decoy sequences were appended using the transproteomic pipeline with Spectrast library generation and conversion to TraML format as described previously^86^. The final library has been provided as a Supplementary file to this report. Peptide identification was performed as previously described^86,87^. MS data and supplementary analysis files were submitted to ProteomeXchange via the PRIDE database^88^ under the identifiers PXD023012 for the dilution series, PXD022996 for the small pilot data set, and PXD023040 for the patient tumor proteomic data set.

### AlphaFold

Wildtype and mutant sequences were obtained from NCBI^89^. AlphaFold server beta^90^ was used to predict protein structure and interactions between wildtype or mutant MDC1 and TP53BP1. The observed ipTM of 0.28 and pTM = 0.3 indicate the prediction was unsuccessful.

## Data Availability Statement

All whole genome and targeted sequencing data will be made publicly available in GEO upon publication of the manuscript. Gene expression (RNA-Seq) and methylation (WGBS) data are publicly available under GSE202245. MS data and supplementary analysis files were submitted to ProteomeXchange via the PRIDE database^88^ under the identifiers PXD023012 for the dilution series, PXD022996 for the small pilot data set, and PXD023040 for the patient tumor proteomic data set.

---

**Supplementary Fig 1.**
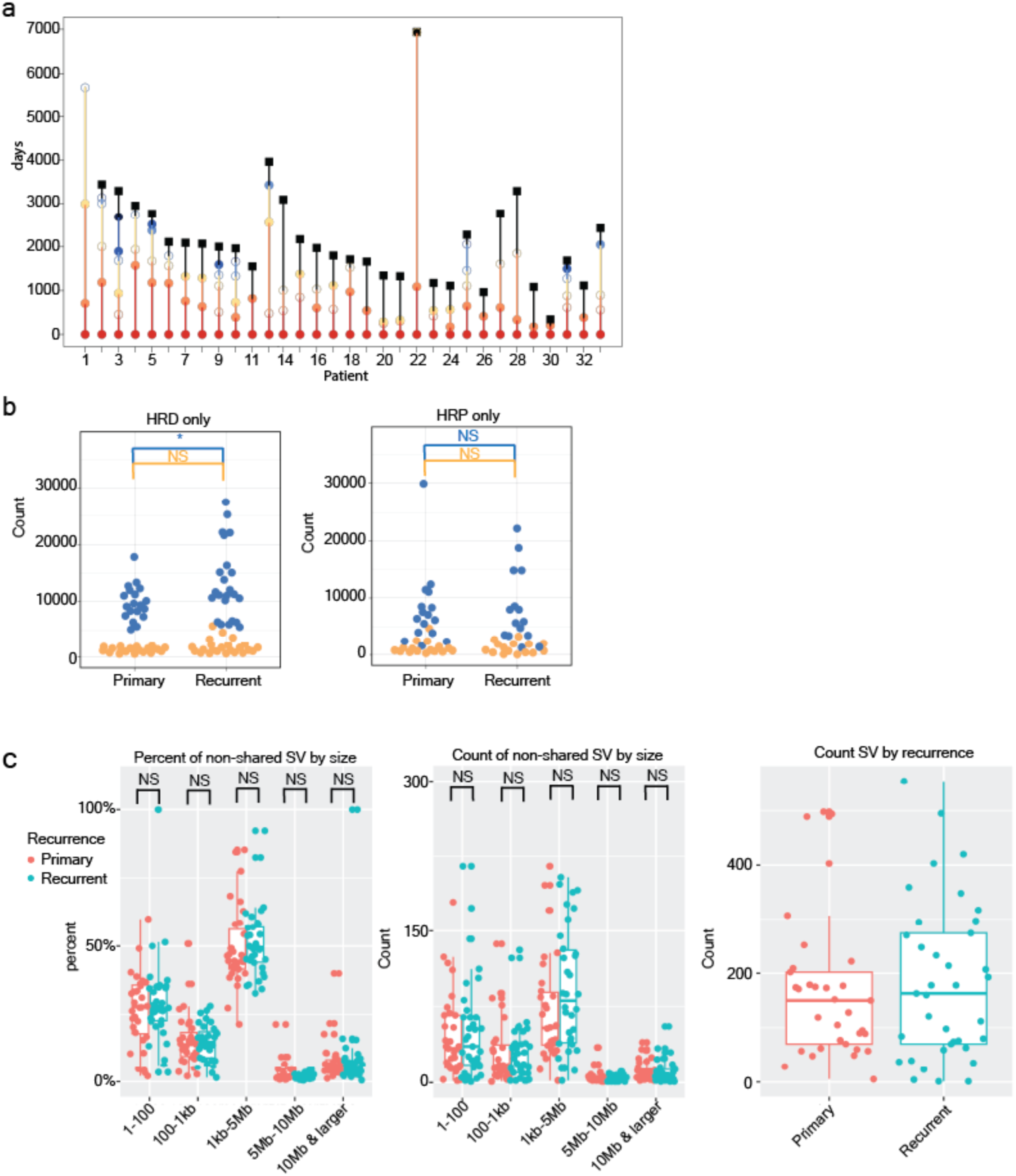
Patient cohort and shared SV. a, Dot plot showing the timeline for each patient. Closed circles are tumors we profiled; open circles are tumors that were resected but not profiled. b, Count of SSM by recurrence, (left) all HRD patients (right) all HRP patients. c, Structural variants binned by size. (Left) percent of non-shared SV per patient, (middle) count of non-shared SV per patient, (right) count of all SV stratified by recurrence. *p < 0.05, NS not significant, students T test.

**Supplementary Fig 2.**
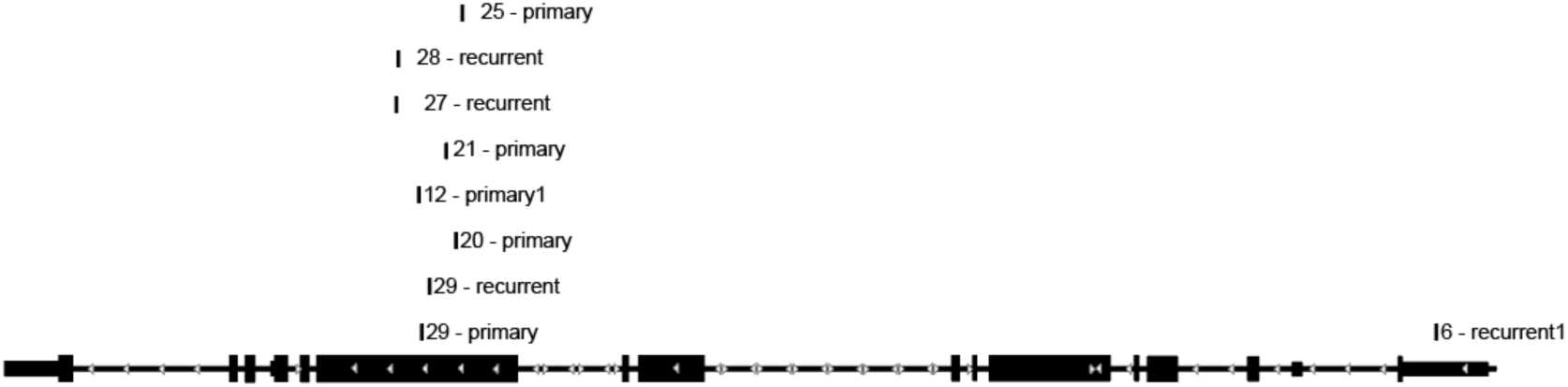
MDC1 mutations are clustered in exon 11. Schematic displaying the location of each MDC1 mutation found in our cohort.

**Supplementary Fig 3.**
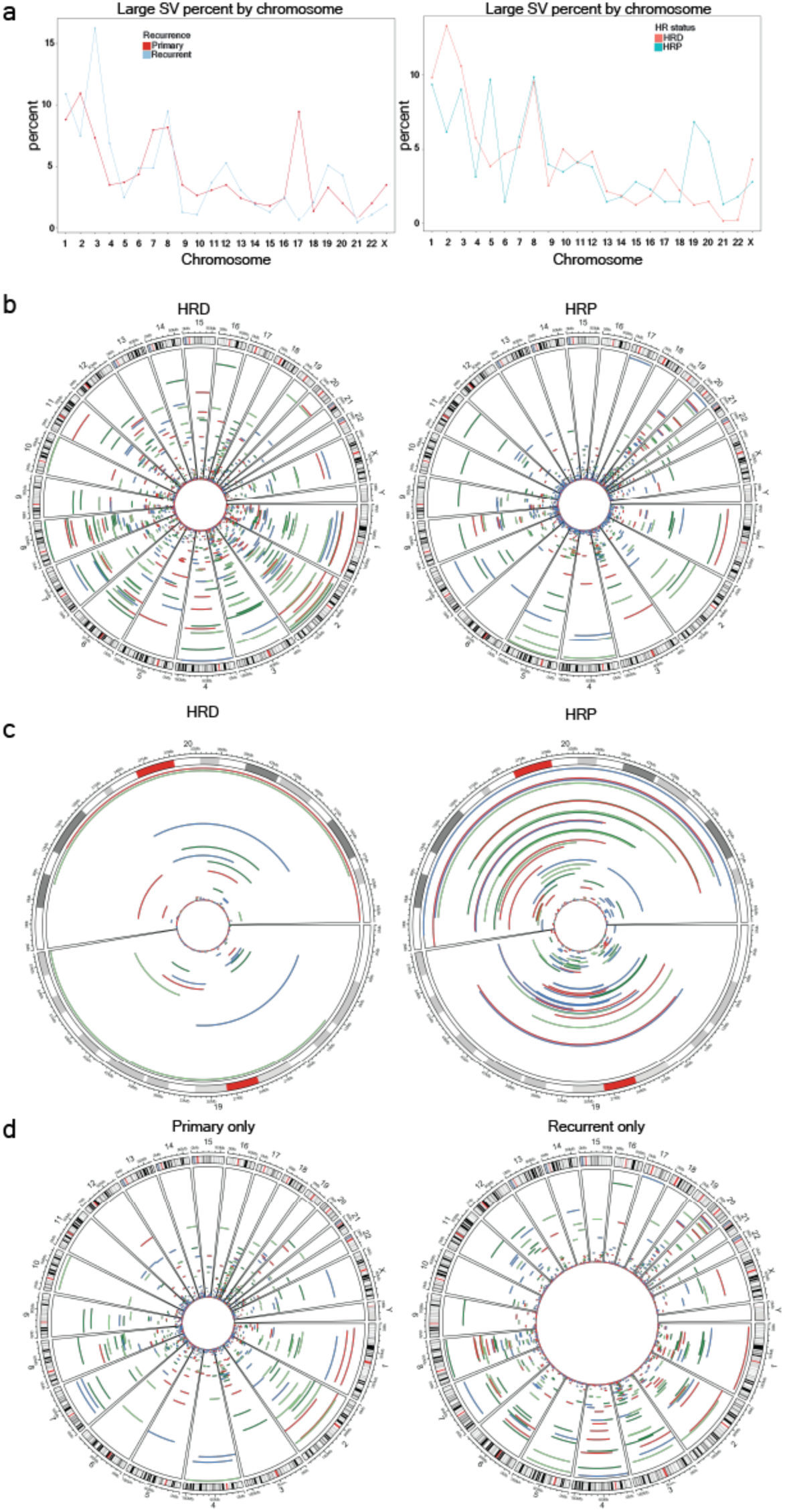
SV size and distributions. a, Percent of large (<5mb) SVs in the genome split by recurrent status (left) and HR status (right). b, All SVs split by HRD and HRP. c, Plots depicting the chromosome 19 and 20 SVs split by HRD and HRP tumors. d, All SV split by primary and recurrent sv status

**Supplementary Fig 4.**
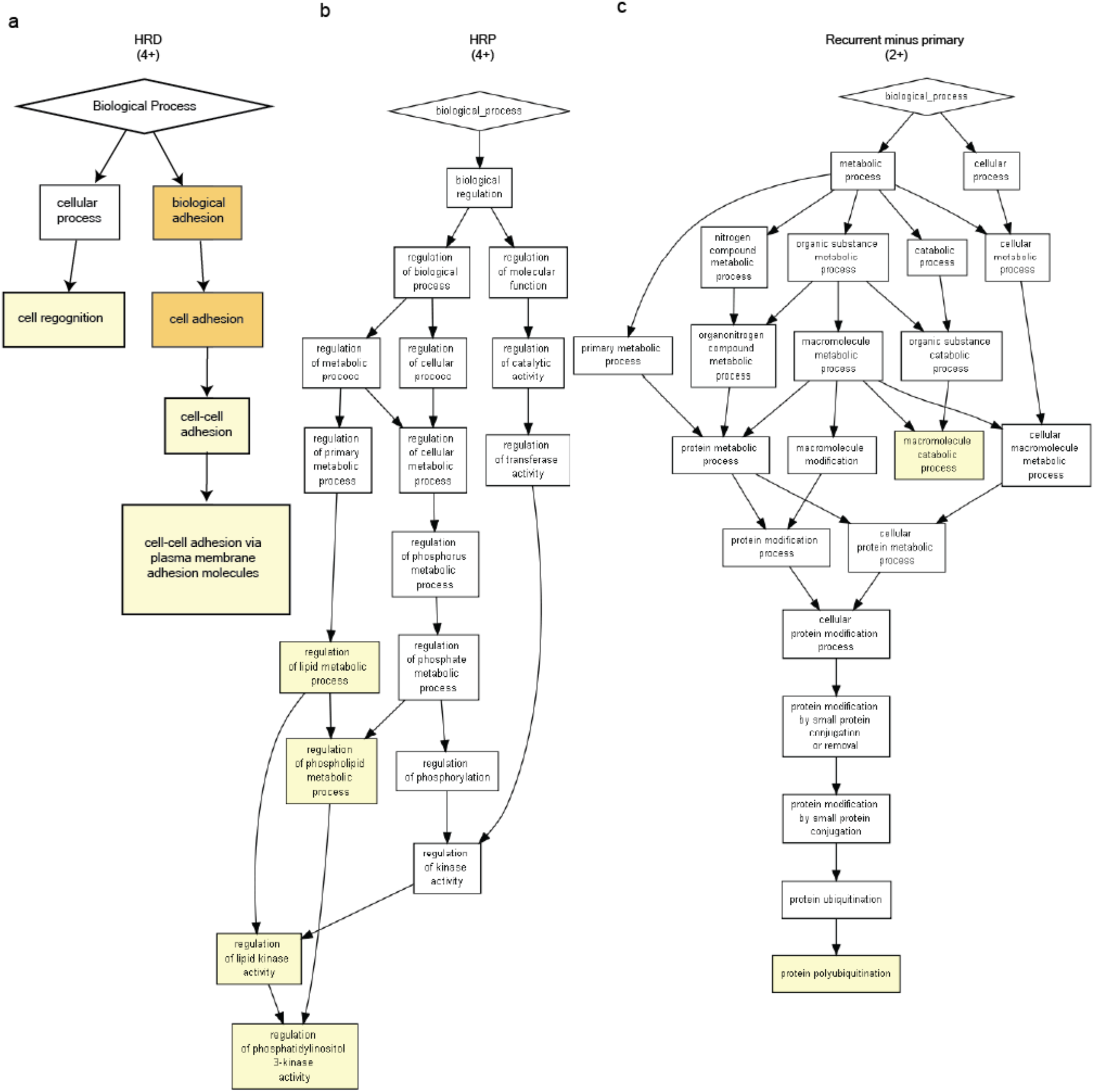
Gene Ontology. a, Gene ontology plot depicting the HRD specific biological processes altered due to deletions, translocations, inversions, and duplications with a breakpoint in the gene body of genes found in at least 4 tumors. b, Gene ontology plot depicting the HRP specific biological processes altered due to deletions, translocations, inversions, and duplications with a breakpoint in the body of genes found in at least 4 tumors. c, Gene ontology plot depicting the recurrent specific biological processes altered due to deletions, translocations, inversions, and duplications with a breakpoint in the body of genes found in at least two tumors. For a-c, p value based on mHG scores. White boxes indicate p>10^−3^, yellow boxes indicate p<10^−3^ & >10^−5^, orange boxes indicate 10^−5^ & p<10^−7^

**Supplementary Fig 5.**
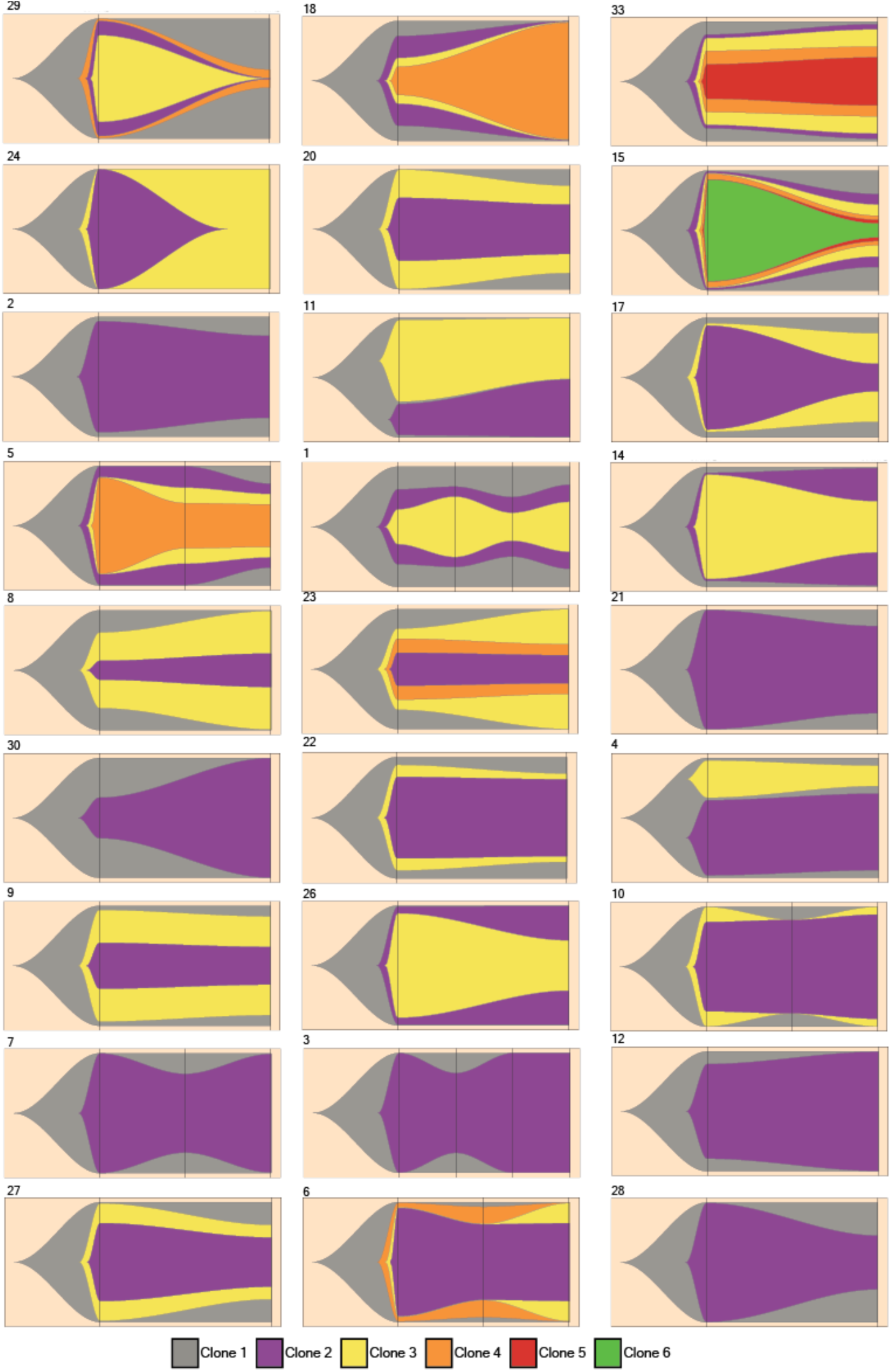
SNV clonal dynamics. Fishplots for each patient in our cohort. Black lines indicate sampling and are evenly spaced irrespective of time between samplings. Clones arising from within another clone are direct descendants of that clone. Each panel is labeled with the Patient number in the top left of the panel.

**Supplementary Fig 6.**
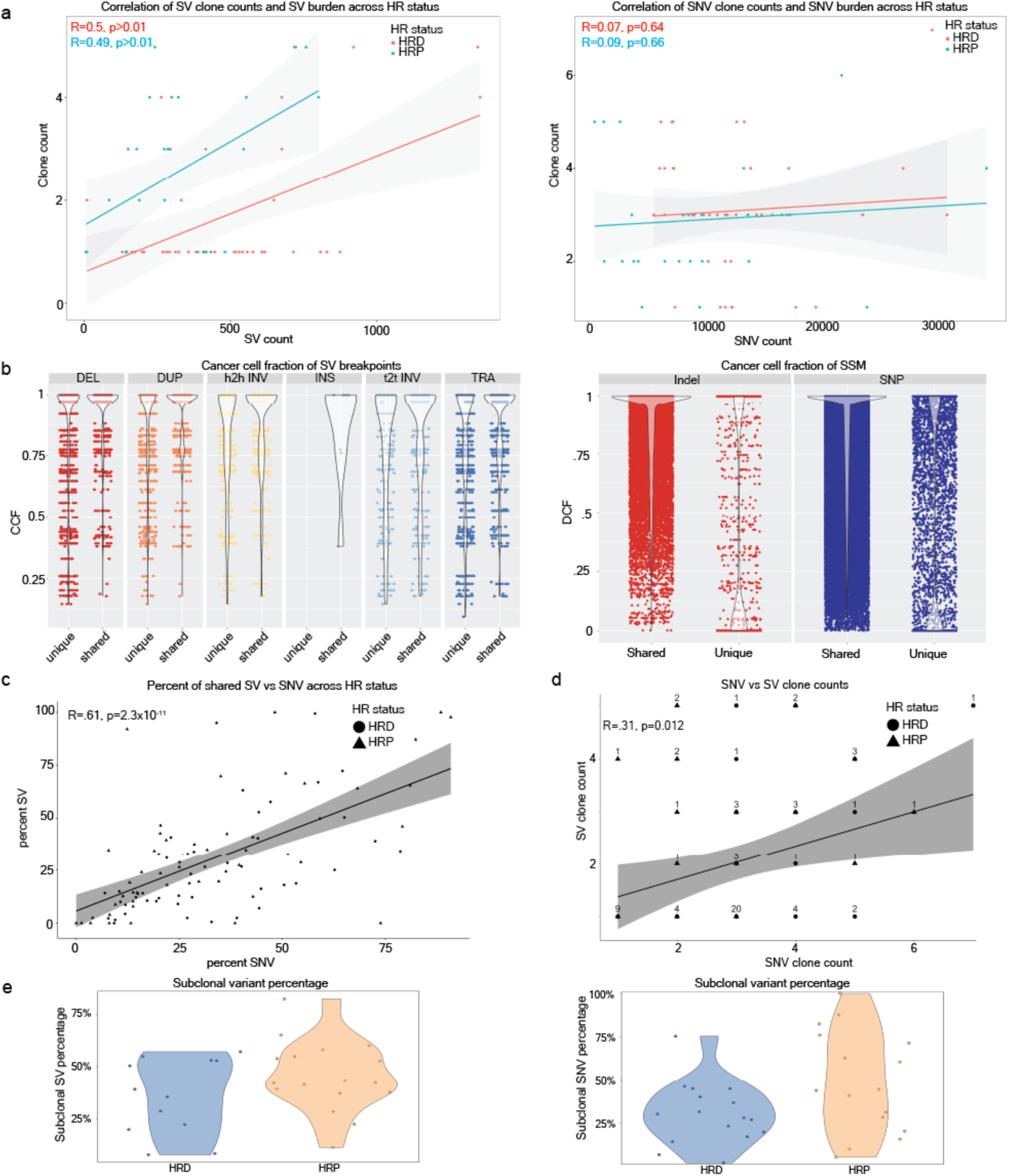
Clonal outcomes of SV and SNV. a, correlation of (left) SV count and (right) SNV counts with clone counts based off of each metric. b, cancer cell fraction for (left) each SV breakpoints or (right) SNV and indels. c, correlation of percent of shared SV versus SNV for each tumor in the cohort. d, correlation of clone counts based on SV or SNV analysis, number above dots indicate the number of overlapping symbols. e, percent of subclonal (left) SV or (right) SNV per tumor, split by HR status.

**Supplementary Fig 7.**
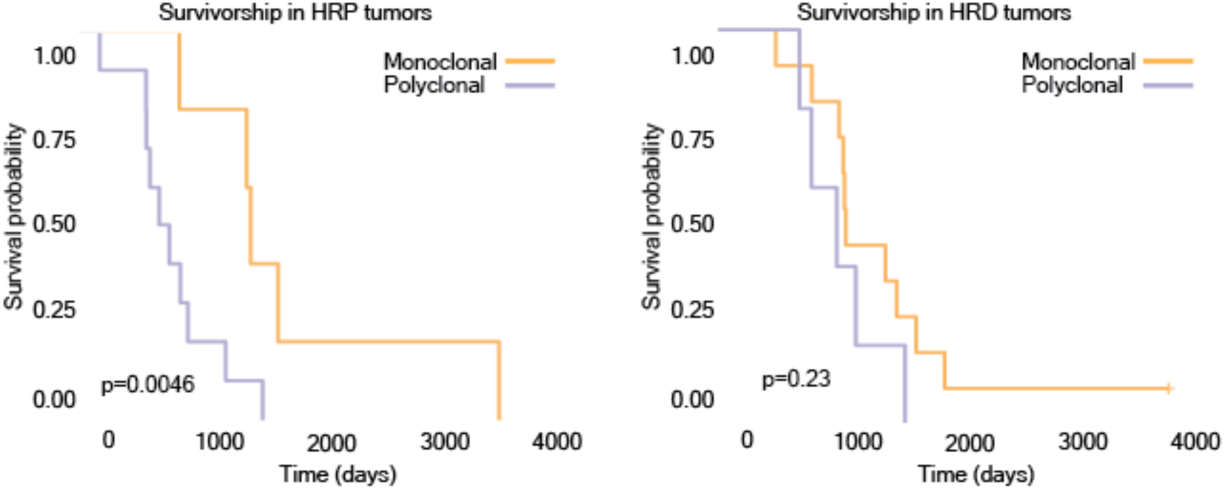
Clonality and SV groups affect patient survival. Kaplan-Meier plot showing the survivorship of HRP (left) or HRD (right) patients with monoclonal or polyclonal primary tumors.

**Supplementary Fig 8.**
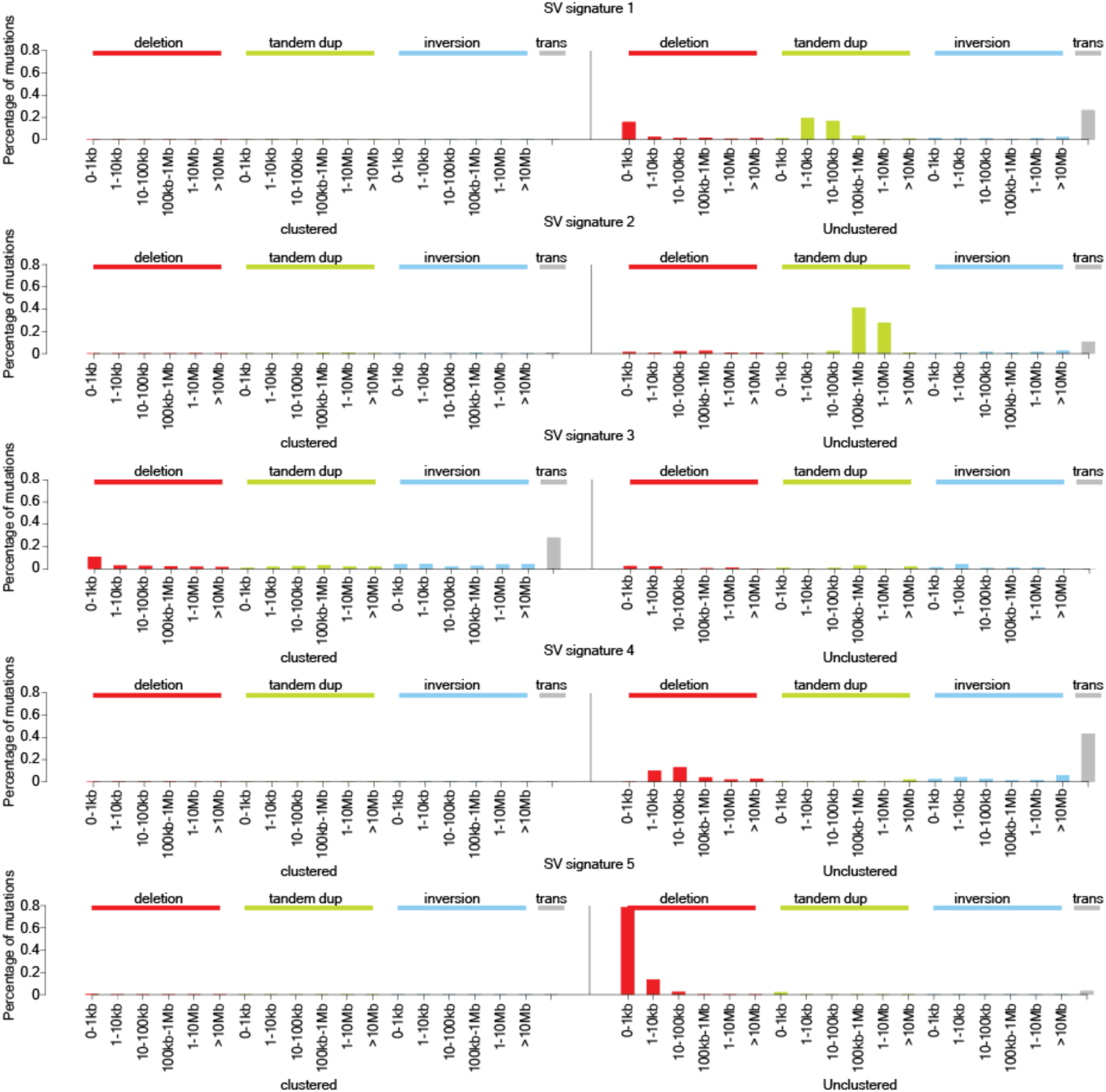
Five *de novo* SV signatures were identified in our cohort. SV signatures displaying the percent of mutations in that signature that were a given SV type within a given size group. SV was determined to be part of a mutation cluster are displayed on the left, while those on the right were unclustered. Clustered translocations are indicative of copy number neutral insertions.

**Supplementary Fig 9.**
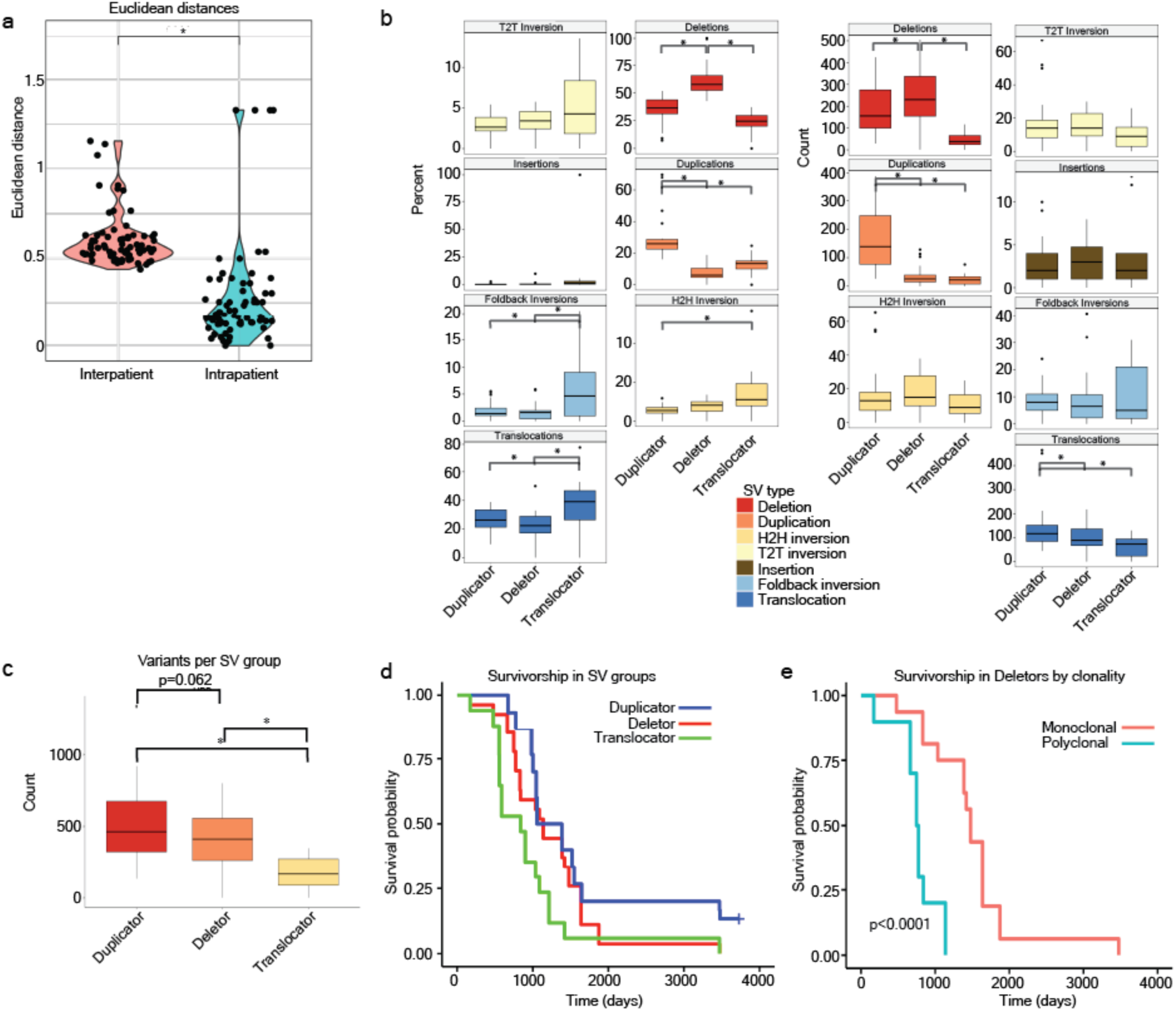
SV groups have differential SV outcomes. a, Euclidean distances between tumors from the same and different patients. b, Percent of SV present in a tumor (left) or total count (right) for each SV type in each SV group. c, Total variants in each SV group. d, Kaplan-Meier plot showing the survivorship of each SV group. e, Kaplan-Meier plot showing the survivorship of deletor SV group patients with monoclonal or polyclonal primary tumors. *p < 0.05, NS not significant, students T test.

**Supplementary Figure 10.**
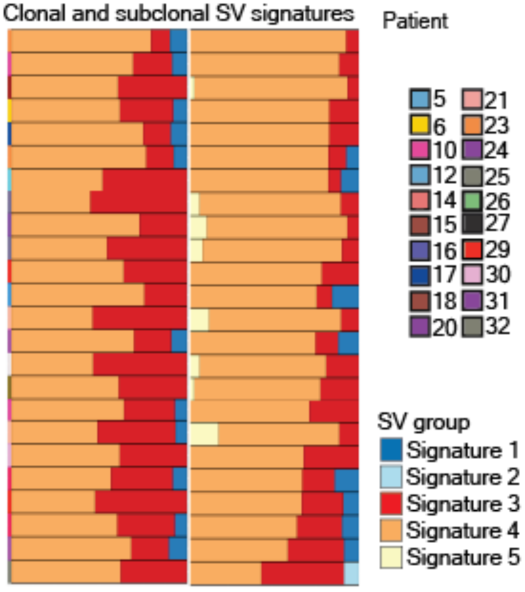
clonal and subclonal SV signatures. Clonal (left) and subclonal (right) SV signatures for each polyclonal tumor in our cohort.

**Supplementary Fig 11.**
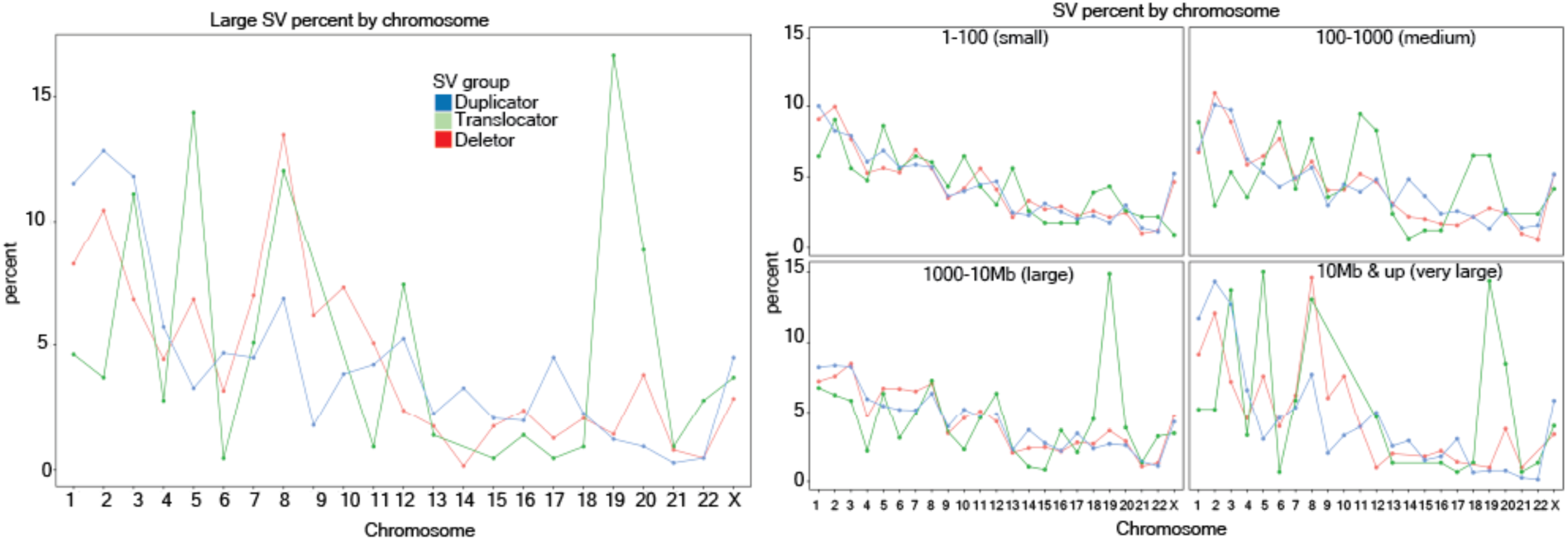
SV size by group. Percent SV in each size group per chromosome, split by group. SVs 5Mb and larger (left) and all other size groups (right)

**Supplementary Fig 12.**
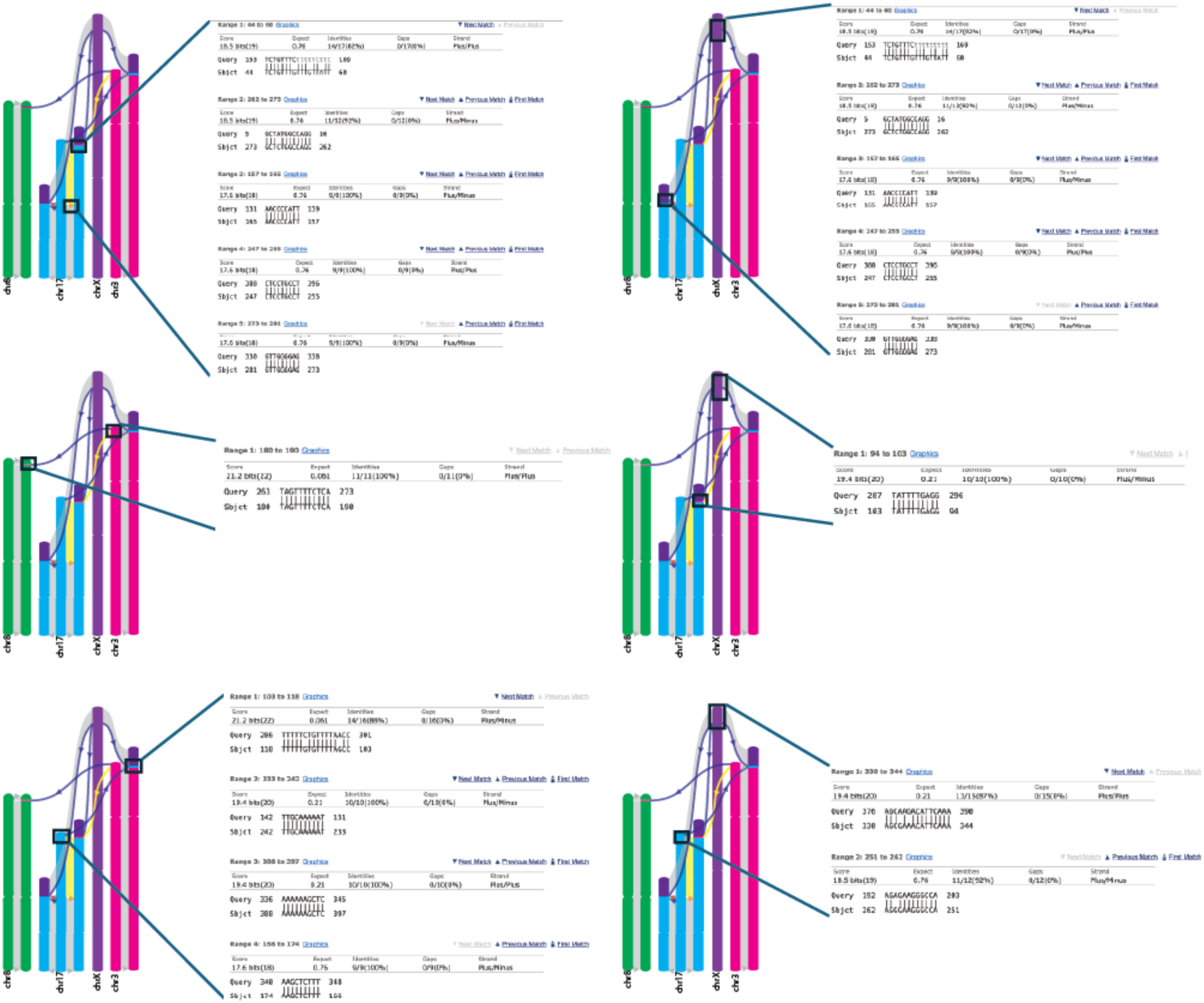
Homology directed repair may influence the creation of cluster 1. Regions around cluster 1 breakpoints in the recurrent tumor of patient 14 were extended 200 base pairs and input into blast. (left) Diagrams with boxes depicting the two blast regions, (right) screenshots of the blast output.

**Supplementary Fig 13.**
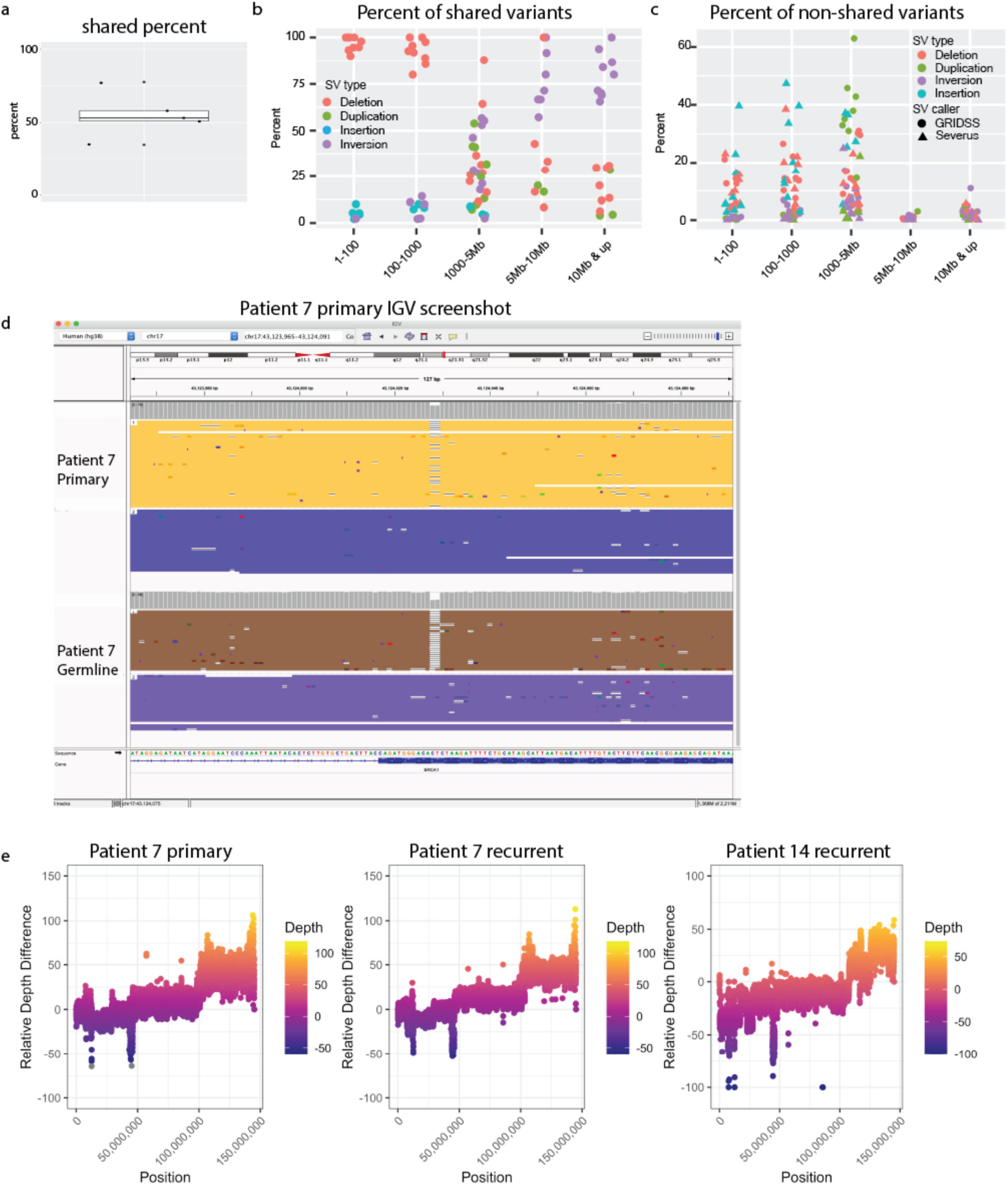
Long read data shares many variants and identifies new variants. a, Percent of shared variants within the caller that identified fewer SV. b, Size and variant type of SV identified in both short read and long read sequencing. c, Percent and type of variants that were unique to each variant caller, binned by SV size. d, IGV screenshot of the *BRCA1* mutation that displays subclonal *BRCA1* reversion patient 7s primary tumor. e, relative depth plots that display the tumor depth /depth of germline for a given region. The region displayed here is chromosome 8, in which all have q arm amplifications leading to large copy number shifts.

## Notes

### Competing Interest Statement

The authors have declared no competing interest.

